# Noradrenergic efferent subsystems that gate traumatic social learning

**DOI:** 10.1101/2025.03.07.641347

**Authors:** Kaoru Seiriki, Shunsuke Maeda, Leo Kojima, Yuzuka Fujimoto, Taiyou Baba, Hiroki Rokujo, Tomoki Nitta, Takanobu Nakazawa, Atsushi Kasai, Takatoshi Hikida, Hitoshi Hashimoto

## Abstract

Individuals experience varying magnitudes of stress in their daily lives. Although stress responses facilitate adaptive processes to cope with changing environments, severe stress can lead to traumatic learning and anxiety disorders. However, the neuronal mechanisms underlying the influence of stress severity on these processes remain unclear. Here, we show that traumatic social stress engages anatomically distinct locus coeruleus noradrenergic (LC^NA^) subpopulations that exhibit dynamic responses scaled to aversive salience. Using whole-brain activity and axonal projection mappings, we identified projectome subtypes of LC^NA^ neurons, which are differentially recruited by single versus consecutive aversive social experiences. While hippocampus-projecting LC^NA^ neurons responded to general social contacts, thalamus-projecting LC^NA^ neurons tracked the aversive salience of social stimuli. Functional manipulations revealed a bidirectional role of thalamus-projecting LC^NA^ neurons in social avoidance learning. These findings reveal the functional architecture of LC^NA^ subsystems that regulate traumatic social learning via dynamic scaling to aversive salience.

## INTRODUCTION

Stress induces various physiological and behavioral responses to cope with real and potential threats, depending on the stress type, duration, and intensity^1–4^. However, severe stress, even from a single event, can be traumatic and may trigger a stress-related psychiatric disorder^5–7^. Clinical studies have suggested that traumas related to interpersonal violence are a strong risk factor for posttraumatic stress disorders^7,8^. Compared to other stress types, social stress in rodents can induce severe behavioral and physiological symptoms depending on the stress severity, including stress frequency and duration^9–11^. Social avoidance and impaired social motivation are critical consequences of severe social stress and are mediated through specific neuronal circuits^12–15^. However, the neurobiological relationships between stress conditions and resulting sociobehavioral plasticity remain unclear.

During stress responses, various neuromodulators, including neuropeptides and monoamines, are released with spatial and temporal variations depending on the stress types and intensities^2^. Although accumulating evidence suggests that the noradrenergic inputs to the amygdala mediate enhanced fear and stress responses^16,17^, the differential effects of these neuromodulators across brain regions are thought to contribute to distinct symptoms of anxiety disorders^18^. Therefore, there is still a need to elucidate how and which neuromodulatory circuitries orchestrate the stressor-specific actions for driving behavioral switching. Recent studies have demonstrated that monoaminergic systems comprise heterogeneous cell subpopulations, which are each characterized by distinct projection targets and specific behavioral roles^19–24^, including anxiety^17,23,25^ and threat learning^22^. Despite the emergence of subpopulation-specific neuronal substrates of general stress-related behaviors, it is important to extend this knowledge to understand whether and how anatomically distinct subpopulations are recruited based on the frequency and duration of stress events. Given the stressor-specific activity patterns across multiple brain region^15^ and the extensive axonal wiring of neuromodulatory systems at a brain-wide scale^23,26–28^, it is essential to perform a mesoscopic characterization of neural response differences across stress conditions and their associated circuit structures. In this study, we identified distinct LC^NA^ circuits that exhibit differential neural responses depending on social stress conditions, leading to social avoidance learning through projections to the thalamus rather than the hippocampal and cortical areas.

## RESULTS

### Brain-wide activation patterns following different types of stress

To characterize neuronal activities across different stress intensities and types, whole-brain mapping of the immediate early gene Fos was performed using Fos-enhanced green fluorescent protein (EGFP) reporter mice^29^ and confocal serial section whole-brain imaging system, block-FAce Serial microscopy Tomography (FAST)^30,31^. For activity mapping, Fos-EGFP mice were subjected to home cage (Naïve), single social defeat stress (SD×1), acute consecutive social defeat stress (SD×5), restraint stress (Res), novel environment exploration (NE), 0.3 mA weak footshock (FS-0.3), or 0.8 mA strong footshock (FS-0.8) episodes within 1 h (Figs. 1A and 1B). We registered the brains to the Allen reference template, Common Coordinate Framework version 3 (CCFv3)^32^; quantified the number of Fos-positive cells across 250 regions; and applied principal component analysis (PCA) to investigate the brain-wide activation patterns (see Materials and Methods). Visualization of mouse positions in the two-dimensional space defined by the first two principal components (PC1 and PC2) revealed a clear separation of activity patterns between social and non-social stress (Fig. 1C). Analysis of inter-individual Euclidean distances in PC space showed that SD×5 induced the largest shift in brainwide activity patterns from the naïve states, with increased frequency of social stress having a larger effect on activity shifts than increased footshock intensity (Fig. 1D). The difference between SD×1 and SD×5 reflected in PC1, with the highest factor loadings observed in the LC (Fig. 1E). Consistent with this, the LC showed a significant increase in Fos-EGFP+ cell counts in response to SD×5 compared to SD×1 among all brain regions (p < 0.05, increase > 2-fold, and effect size > 3; Figs. 1F, S1A, and S1B). Voxel-based comparison between SD×1 and SD×5 also revealed a significant difference in Fos-EGFP+ cell density in the LC area (Figs. 1G and 1H). Compared with the other stress conditions, SD×5 induced the highest Fos-EGFP+ cell density in the LC (Fig. 1I).

**Fig. 1.**
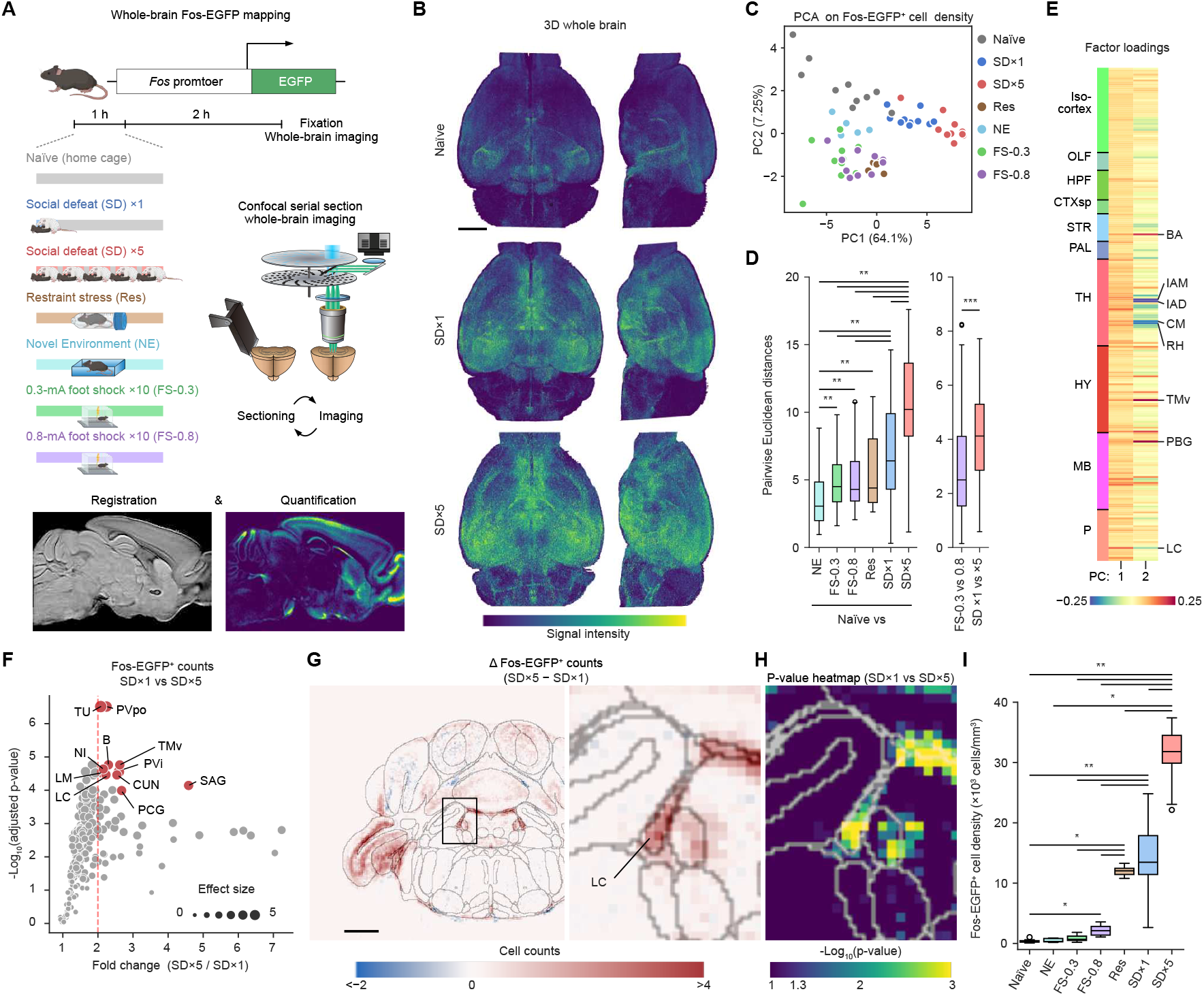
Distinct brain-wide activation signatures across diverse stress conditions. (**A**) Schematic illustrating whole-brain Fos-EGFP mapping. Fos-EGFP mice were subjected to Naïve, SD×1, SD×5, Res, NE, FS-0.3, FS-0.8 conditions and fixed 3 h after the onset of stress conditions. (**B**) Representative images of the three-dimensional (3D) reconstruction of the whole brain. Scale bar, 2 mm. (**C**) PCA of Fos-EGFP+ cell density. Individual mice were plotted on the two-dimensional space defined by the first two principal components (PC1 and PC2). Naïve, n = 10; SD ×1, n = 10; SD ×5, n = 10; NE, n = 5; Res, n = 5, FS-0.3, n = 10; FS-0.8, n = 10. (**D**) Pairwise Euclidean distances in the PC space defined by PC1 and PC2. Comparison of pairwise distances relative to the Naïve group (left) and intra-stress modality distances between social defeat stress and footshock stress (right). **P < 0.01, ***P < 0.001; Kruskal-Wallis test followed by Steel-Dwass test (left) and Mann-Whitney U test (right). (**E**) Heatmap of PCA factor loadings for brain regions. The names of brain regions with higher contributions are indicated. OLF, olfactory areas; HPF, hippocampal formation; CTXsp, cortical subplate; STR, striatum; PAL, pallidum; TH, thalamus; HY, hypothalamus; MB, midbrain; P, pons; BA, bed nucleus of the accessory olfactory tract; IAM, interanteromedial nucleus of the thalamus; IAD, interanterodorsal nucleus of the thalamus; CM, central medial nucleus of the thalamus; RH, rhomboid nucleus; TMv, tuberomammillary nucleus, ventral part; PBG, parabigeminal nucleus. (**F**) Volcano plot showing the adjusted P-value and fold change (SD ×5 over SD×1) of brain regions in mice subjected to social defeat stress. Brain regions with a higher effect size (Cohens d > 3) and fold change > 2 were highlighted with red color. PVi, periventricular hypothalamic nucleus, intermediate part; TU, tuberal nucleus; LM, lateral mammillary nucleus; CUN, cuneiform nucleus; SAG, nucleus sagulum; PCG, pontine central gray; B, Barringtons nucleus; NI, nucleus incertus. (**G**) Coronal view of the voxel heatmap of difference in mean Fos-EGFP+ cell densities (SD×5 − SD×1). The magnified area including the LC was outlined with a black line. Scale bar, 1 mm. (**H**) Magnified coronal view of the voxel heatmap of P-values for the statistical comparison between SD×1 and SD×5. (**I**) Quantification of Fos-EGFP density in the LC induced by different stress types and magnitudes. *P < 0.05, **P < 0.01; Kruskal-Wallis test followed by Steel-Dwass test. For boxplots, boxes indicate the interquartile range with median lines, whiskers extend to 1.5 times the interquartile range, and points represent outliers beyond these ranges. The illustrations of mice in (A) were created with BioRender.

LC neurons are a major source of NA throughout the brain, and are characterized by NA biosynthetic enzymes, including tyrosine hydroxylase (TH)^33^. To examine Fos expression in LC^NA^ neurons, we performed immunohistochemical analysis and found that SD×5 elicited Fos expression in 73% of TH+ neurons in the LC, which was significantly higher than the 36% induced by SD×1 (Fig. S1C). Although the role of the LC^NA^ system in stress responses has been extensively investigated^1,34,35^, our findings suggest that the distinct activity in the LC^NA^ neurons may produce different responses depending on prolonged aversive social events.

### Distinct social behaviors and LC^NA^ activity by stress

To investigate behavioral effects of SD×1 and SD×5, social approach and avoidance behaviors were assessed using open field arena with a social target cage^12,15^ (Fig. 2A). When mice were subjected to SD×1 and SD×5, the SD×5 mouse group showed reduced time in the social zone with an unfamiliar CD1 mouse, increased time in the corner zone, and decreased social approach-avoidance balances compared to both the SD×1 and stress-naïve groups (Figs. 2B–2D). The persistent anxiety-related behaviors days after stress exposure^12^ were not observed in these stress paradigms (Figs. S2A–S2C). To determine the subthreshold level of social defeat stress, we examined shortened versions of SD×2 and SD×3 (see Materials and Methods for details) and found that SD×3, but not by SD×2, induced a reduced preference for the social zone (Figs. 2D–2G).

**Fig. 2.**
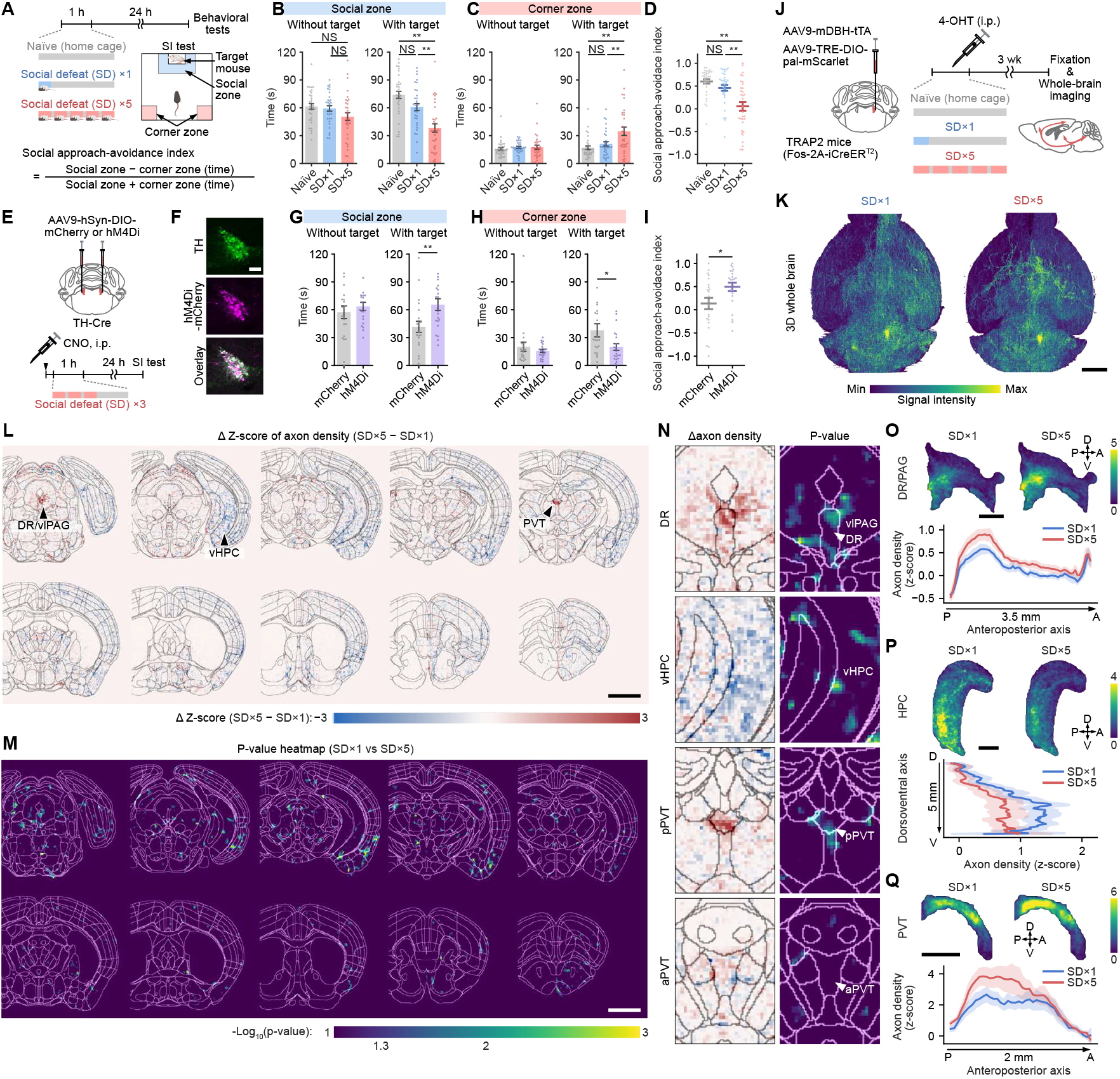
Defeat-induced social avoidance learning and mesoscale projectome of LC^NA^ neurons activated by social defeat stress. (**A**) Schematic illustrating behavioral experiments. Mice underwent social defeat stress episodes followed by a social interaction test at 1 day after stress exposure. (**B, C**) Quantification of time spent in the social and corner zones with or without a target mouse. Naïve, n = 34; SD×1, n = 33; SD×5, n = 33. NS, not significant; **P < 0.01, one-way analysis of variance (ANOVA) followed by Tukeys test. (**D**) Quantification of the social approach avoidance index. **P < 0.01, one-way ANOVA followed by Holm-Sidak test. (**E**) Schematic illustrating behavioral experiments for chemogenetic inhibition of LC^NA^ neurons. Mice were subjected to SD×3 episodes at 15 min after intraperitoneal (i.p.) administration of 5 mg/kg CNO. (**F**) Representative immunofluorescence images of the TH+ neurons expressing hM4Di-mCherry. Scale bar, 100 µm. (**G, H**) Quantification of time spent in the social and corner zones with or without a target mouse in the chemogenetic experiment. mCherry, n = 22; hM4Di, n = 22. *P < 0.05, **P < 0.01, Students t-test. (**I**)Quantification of the social approach avoidance index in the chemogenetic experiment. (**J**) Schematic for mapping whole-brain projection of LC^NA^ neurons activated by different social defeat stress paradigms using TRAP2 (Fos-2A-iCreER^T2^) mice and AAV vectors. (**K**) Representative 3D reconstructions of the whole brain from mice TRAPed by SD×1 or SD×5. Scale bar, 2 mm. (**L, M**) Coronal views of the voxel heatmaps of the difference in z-score-normalized axon densities (SD×5 − SD×1) and P-values for statistical comparison of z-score-normalized axon densities between SD×1 and SD×5. SD×1, n = 5; SD×5, n = 6. DR, dorsal raphe; vlPAG, ventrolateral periaqueductal gray; vHPC, ventral hippocampus; PVT, paraventricular thalamic nucleus. Scale bars, 2 mm. (**N**) Magnified difference heatmaps (left column) and P-value heatmaps (right column) for the DR, vHPC, posterior PVT (pPVT), and anterior PVT (aPVT). (**O–Q**) Heatmap representations of local axon density distribution in lateral view and their quantification along anteroposterior axis for the DR/PAG and PVT (O, Q) or dorsoventral axis for the vHPC (P). Scale bars, 1 mm. Data are represented as mean ± SEM. The illustrations of mice in (A) were created with BioRender.

Next, we investigated the roles of LC^NA^ activation in social trauma-related avoidance learning using Designer Receptors Exclusively Activated by Designer Drugs (DREADD)^36^. Adeno-associated virus (AAV) vectors carrying Cre-dependent inhibitory DREADD hM4Di-mCherry expression cassette were injected into the LC of TH-Cre mice^37^ (Figs. 2E and 2F). Mice expressing hM4Di in the LC were administered intraperitoneal injections of the DREADD ligand Clozapine N-oxide (CNO) at a dose of 5 mg/kg, followed by exposure to SD×3. Chemogenetic inhibition of LC^NA^ neurons during stress episodes resulted in increased time spent in the social zone, reduced time spent in the corner zone, and a social approach-shifted behavioral balance (Figs. 2G–2I). Pharmacological blockade of beta-adrenergic receptor, but not alpha-1-adrenergic receptor, during stress episodes reduced the time spent in the corner zone and increased the social approach-avoidance index (Figs. S2H–2K). These findings demonstrated that the enhanced noradrenergic activity mediates social avoidance learning induced by severe social stress through consecutive defeat episodes.

Given the anatomically and functionally heterogeneous organization of the LC^NA^ system^22,24,25,38^, a specific LC^NA^ circuitry might mediate social trauma-related avoidance learning. To determine the circuit structures of the LC^NA^ neuronal ensembles recruited by single and consecutive social defeat stress episodes, we employed intersectional approaches using the Cre recombinase and Tet-Off system. First, we constructed an AAV vector to specifically express fluorescent proteins in the LC^NA^ neurons using a mouse dopamine beta-hydroxylase (mDBH) promoter with our Tet-Off system-based cell type-specific AAV approach, which is named TransActivator-Regulated Enhanced Gene Expression within Targeted neuronal population (TAREGET)^39^ (Fig. S3A). The TAREGET AAV (AAV9-mDBH-tTA) allowed a 92% specificity for fluorescent protein expression in LC^NA^ neurons under the control of the tetracycline-responsive element promoter (TRE) (Figs. S3B and S3C). A combination of AAV9-mDBH-tTA with AAV9-TRE-pal-mScarlet^39^, which is the membrane-targeted red fluorescent protein, facilitated brain-wide labeling of LC^NA^ axons for whole-brain imaging (Figs. S3D–S3F). Using these strategies, an entire structure of LC^NA^ circuitry was identified, and there were no sex differences throughout the brain (Fig. S3G).

For activity-dependent labeling, we employed targeted recombination in active populations (TRAP2) mice^40^, where tamoxifen-inducible Cre recombinase (CreER^T2^) is co-expressed with endogenous Fos. To TRAP activated LC^NA^ neurons, TRAP2 mice that received injections of AAV9-mDBH-tTA and AAV9-TRE-DIO-pal-mScarlet into the LC were subjected to home cage, SD×1, or SD×5 conditions, followed by the administration of an active metabolite of tamoxifen, 4-hydroxy tamoxifen (4-OHT) (Fig. 2J). In the stress-naïve group, a small number of TRAPed neurons expressing pal-mScarlet were observed, and the fluorescence of their axons was sparse and dim (Figs. S4A and 4B). Contrastingly, LC^NA^ neurons TRAPed by SD×1 and SD×5 episodes exhibited broadly distributed fluorescence-labeled axons in a three-dimensional (3D) space (Fig. 2K). Although region-based comparison using the parcellated Allen CCFv3 did not reveal significant differences in axon density among various brain regions (Fig. S4C), voxel-based subregional comparison identified substructures that received differential proportions of axonal innervations (Figs. 2L–2N and S4D). The axonal distribution of SD×1-TRAPed neurons was weighted in the ventral hippocampus (vHPC), whereas axonal projections from SD×5-TRAPed neurons were preferentially distributed toward the posterior paraventricular thalamic nucleus (PVT), dorsal raphe (DR), and ventrolateral periaqueductal gray (vlPAG) (Figs. 2N–2Q and S4D). Taken together, these results suggest that consecutive social stress episodes activate anatomically distinct subsets of LC^NA^ neurons to facilitate social avoidance learning.

### Mesoscale projectome of anatomically defined LC^NA^ subsystems

Given the complex wiring patterns of LC^NA^ neurons, including selective projections to specific brain regions^22^, divergent projections to multiple areas^41^, or combinations of both^24,26,28^, it is important to investigate the mesoscale architecture of LC^NA^ projection patterns. To map the brain-wide axon collaterals of LC^NA^ neurons projecting to the vHPC, PVT, or DR (LC^NA^→vHPC, LC^NA^→PVT, and LC^NA^→DR, respectively), each cell population was retrogradely targeted by an AAV2retro vector^42^ encoding Cre (AAVrg-hSyn-iCre). Subsequently, their axonal distribution was analyzed by whole-brain imaging (Fig. 3A). Quantification of labeled cell bodies demonstrated that the LC^NA^→vHPC neurons were characterized by a greater number of cells and broader distribution across the anteroposterior axis, compared to the other two populations (Figs. S5A–5C). Local magnification and mapping the mean axon density in the coronal two-dimensional space showed that LC^NA^ neurons retrogradely targeted from the PVT exhibited axonal innervations to the DR, and vice versa (Figs. 3B and 3C), suggesting the existence of subpopulations projecting to both the PVT and DR. Cluster analysis of axon densities across brain regions revealed that the LC^NA^→vHPC neurons exhibited a widespread projection to the hippocampus and other limbic-related areas (e.g., entorhinal cortex, basolateral amygdala), which was distinct with the LC^NA^→PVT and LC^NA^→DR neurons (Fig. 3D). In contrast, the LC^NA^→PVT and LC^NA^→DR neurons exhibited similar projection patterns at a brain-wide scale (Figs. 3D, 3E, and S5D). However, at a local scale, different projection patterns were observed between the LC^NA^→PVT and LC^NA^→DR neurons, such that LC^NA^→PVT neurons exhibited more innervations to the mediodorsal and midline thalamus (Figs. S5E and S6). Axon densities in the targeted areas (vHPC, PVT, or DR) exhibited heterogeneous distribution patterns across each area (Fig. 3F). We found that the LC^NA^→vHPC neurons project widely throughout the hippocampal area, with significantly higher axon density in the ventral and intermediate portions of the hippocampus (Fig. 3G). Similarly, the LC^NA^→PVT and LC^NA^→DR neurons showed denser projections in the posterior part compared to the anterior part of the PVT and DR, respectively (Figs. 3G and S6). These results suggest that the LC^NA^ neurons contain highly collateralized subsystems, each characterized by distinct projection patterns, including dense collaterals to the hippocampus and its related cerebral regions as well as to the diencephalic and brainstem regions.

**Fig. 3.**
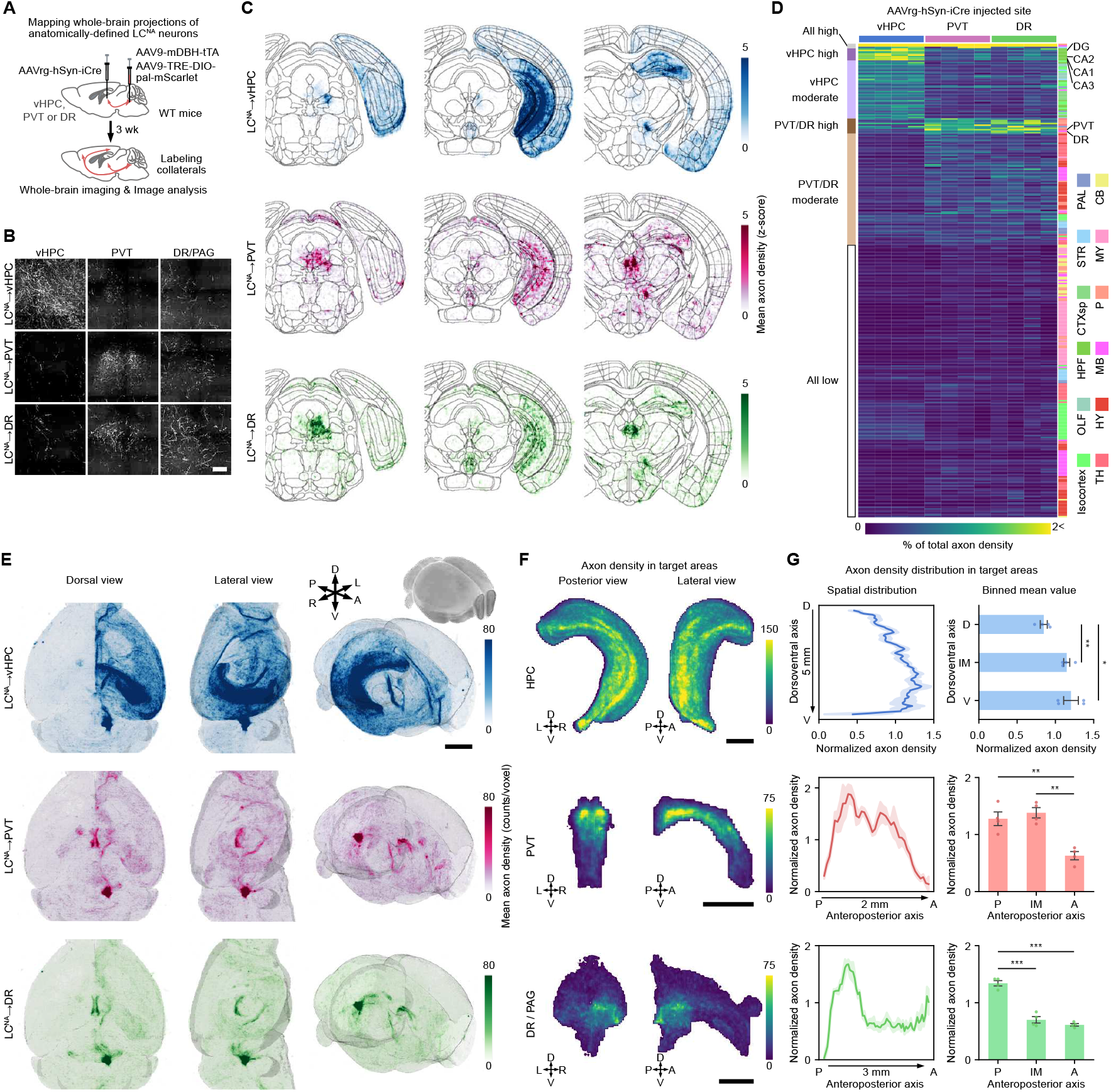
Distinct projectome subtypes in the LC^NA^→vHPC, LC^NA^→PVT, and LC^NA^→DR neurons. (**A**) Schematic for mapping brain-wide collateralization of LC^NA^ neurons projecting to the vHPC, PVT, or DR. (**B**) Representative images of the vHPC, PVT, and DR from mice that underwent projection target-selective labeling. Scale bar, 100 µm. (**C**) Coronal views of the voxel heatmap of z-score-normalized axonal densities. (**D**) Clustered heatmap of the relative axon density from individual mice, with adjustment based on the volume of each brain region and the total label densities across 311 brain regions. The brain regions were clustered using the k-means algorithm. The left color labels indicate the cluster assignments; all high, vHPC high, vHPC moderate, PVT/DR high, PVT/DR moderate, and all low clusters. The right color labels indicate anatomical division based on Allen atlas CCFv3. (**E**) Whole-brain 3D views of the voxel heatmap of the mean axon density. A, anterior; P posterior; L, left; R, right; D, dorsal; V, ventral. Scale bar, 2 mm. (**F**) Heatmap representation of local axon density distribution in the HPC of LC^NA^→vHPC neurons (dop), PVT of LC^NA^→PVT neurons (middle), and DR/PAG of LC^NA^→DR neurons (bottom). (**G**) Quantification of mean axon densities along dorsoventral (HPC) and anteroposterior (PVT, DR/PAG) axes. The distribution was divided into three bins for statistical comparison. D, dorsal; V, ventral; P, posterior; A, anterior; IM, intermediate. *P < 0.05, **P < 0.01, ***P < 0.001, one-way repeated measures ANOVA followed by Holm-Sidak test. Data are represented as mean *±* SEM.

### Neural reactivity of anatomically defined LC^NA^ subsystems

A previous *in vivo* electrophysiological study showed that LC^NA^ neurons can have both shared and distinct response properties depending on the type of environmental stimuli, i.e., a global response to aversive noxious stimuli and a sub-populational response to sensory predictive cue associated with aversive stimuli^22^. To elucidate the relationship between neural response variability and anatomical heterogeneity, we performed fiber photometry recording of neural activity in anatomically defined LC^NA^ subpopulations during social stress episodes. The LC^NA^→vHPC, LC^NA^→PVT, and LC^NA^→DR subpopulations were retrogradely targeted to express genetically encoded Ca^2+^ indicator jG-CaMP7s^43^ (Figs. 4A and S7A). We divided a recording session into three phases: a baseline phase in an aggressors home cage without the aggressor, a sensory contact phase with the aggressor mouse placed in a grid cage, and a physical contact phase with the freely behaving aggressor mouse (Figs. 4B, S7B, and S7C). Subject mice received these sessions 5 times with different aggressors, and the number of attacks during the physical contact phase was controlled using a cage divider to ensure comparable levels of defeat across subjects (Fig. S7D; see Materials and Methods for details). The investigation behavior during sensory contact phases declined across repeated sessions (Fig. S7E). Therefore, we analyzed neural responses to the first five investigations across the first three sessions. We found that social investigation during the unstressed first session elicited a significant increase of the Ca^2+^ responses in the LC^NA^→vHPC, but not LC^NA^→PVT or LC^NA^→DR neurons (Figs. 4C–4E), suggesting that LC^NA^→vHPC neurons are responsive to general social stimuli. In contrast, the LC^NA^→PVT and LC^NA^→DR neurons exhibited significant neural responses to social stimuli after repeated stress experiences. Notably, repeated stress sessions progressively enhanced neural responses in LC^NA^→PVT neurons, not only during social investigation but also at the onset of the social contact phase (Figs. 4D, S7F, and S7G). Consistent with the global response of LC^NA^ neurons to painful stimuli reported in the previous study^22^, attacks by the aggressor accompanied by aversive, noxious stimuli elicited comparable Ca^2+^ responses across all three populations (Figs. 4F–4H). Taken together, these findings indicate that anatomically distinct LC^NA^ subpopulations exhibit differential response properties to salient stimuli during aversive social experiences.

**Fig. 4.**
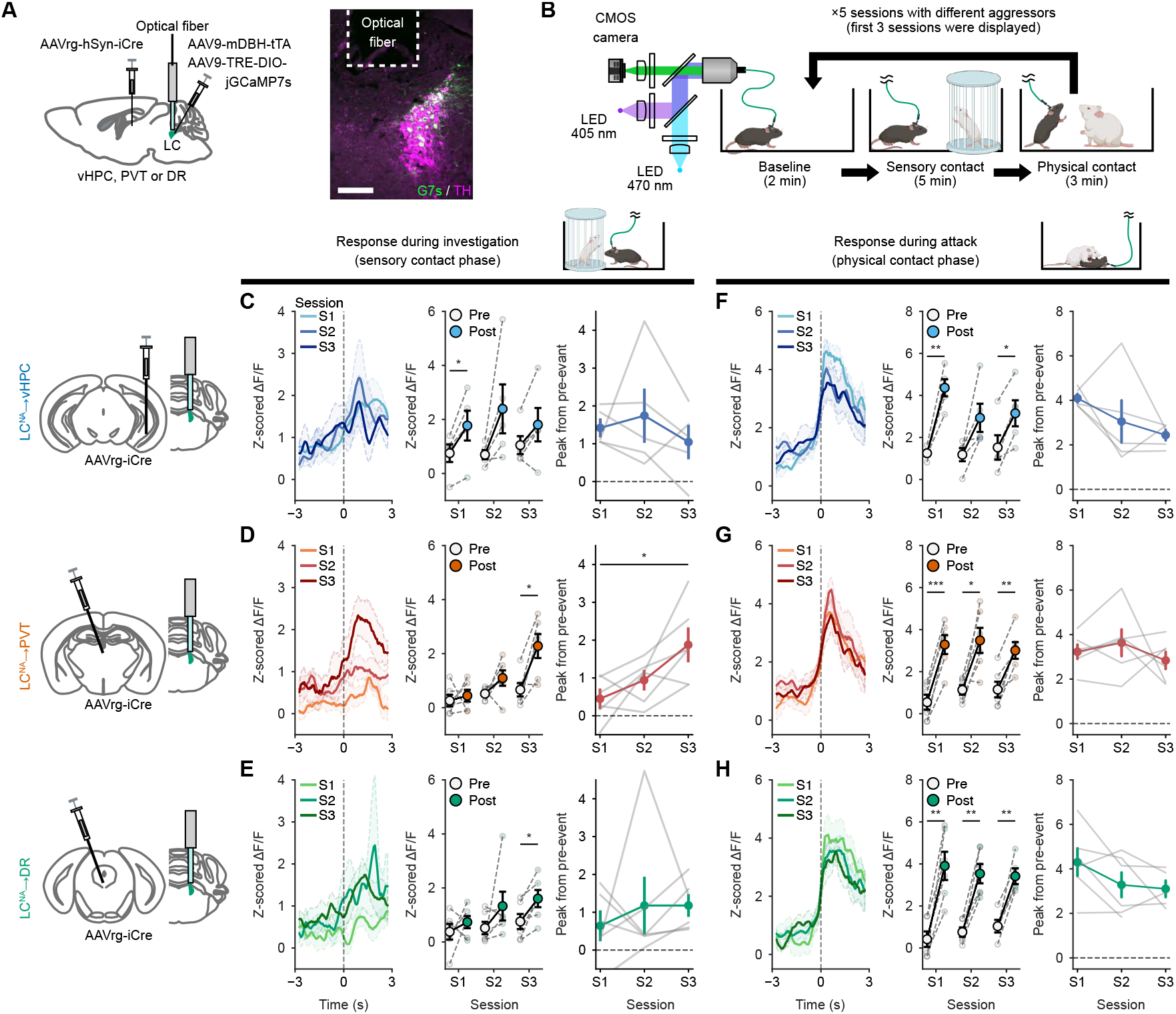
Distinct neural coding of salience of social stimuli in LC^NA^ efferent subsystems. Schematic illustrating the projection target-specific and cell type-selective expression strategy for jGCaMP7s (G7s). The representative image shows jGCaMP7s-expressing LC^NA^ neurons retrogradely targeted from the vHPC. Scale bar, 200 µm. (**B**) Schematic illustrating the fiber photometry recording of jGCaMP7s fluorescence during social defeat stress episodes. A single stress session consists of baseline (2 min), sensory contact (5 min), and physical contact (3 min) phases; further, it was repeated five times. (**C–E**) Neuronal activities of the LC^NA^→vHPC (C), LC^NA^→PVT (D), and LC^NA^→DR (E) neurons during social investigation in the sensory contact phase across sessions (S1–3). Peri-event plot showing mean neuronal activity 3 s before and after social approach (left). Quantification of mean neuronal activity before and after the event onset (middle) and peak response amplitudes after the event onset (0 to 1.5 s) from pre-event baseline (right). (**F–H**) Neuronal activities of the LC^NA^→vHPC (F), LC^NA^→PVT (G), and LC^NA^→DR (H) neurons in response to aggressive attacks during the physical contact phase across sessions. Peri-event plot showing mean neuronal activity 3 s before and after aggressive attack (left). Quantification of mean neuronal activity before and after the event onset (middle) and peak response amplitudes after the event onset (0 to 1.5 s) from pre-event baseline (right). LC^NA^→vHPC, n = 5; LC^NA^→PVT, n = 6; LC^NA^→DR, n = 6. *P < 0.05, **P < 0.01, two-way repeated measures ANOVA followed by pared t-test with Bonferroni correction (pre- and post-event comparison) or one-way repeated measures ANOVA followed by Holm-Sidak test (peak response amplitudes). Data are represented as mean *±* SEM. The illustrations of mice in (B) were created with BioRender.

### Role of LC^NA^ subsystems in social avoidance learning

Finally, we examined whether manipulation of anatomically defined LC^NA^ neurons would modulate social avoidance learning. We first found sparse expression of DREADD hM4Di when controlled using AAV9-mDBH-tTA and constructed a modified tTA comprising the TetR and p65 activation domain (TetR-p65AD) to enhance the expression level^44^ (Fig. S8A). Using these optimized expression strategies, we tested whether chemogenetic silencing of LC^NA^ subsystems during stress experiences affects social avoidance learning (Figs. 5A and S8B). Inhibition of LC^NA^→vHPC neurons during SD×3 episodes did not affect social investigation and avoidance behaviors in the social interaction test, whereas silencing LC^NA^→PVT or LC^NA^→DR neurons significantly increased the time for social investigation as well as the social approach-avoidance balance (Figs. 5B–5D).

**Fig. 5.**
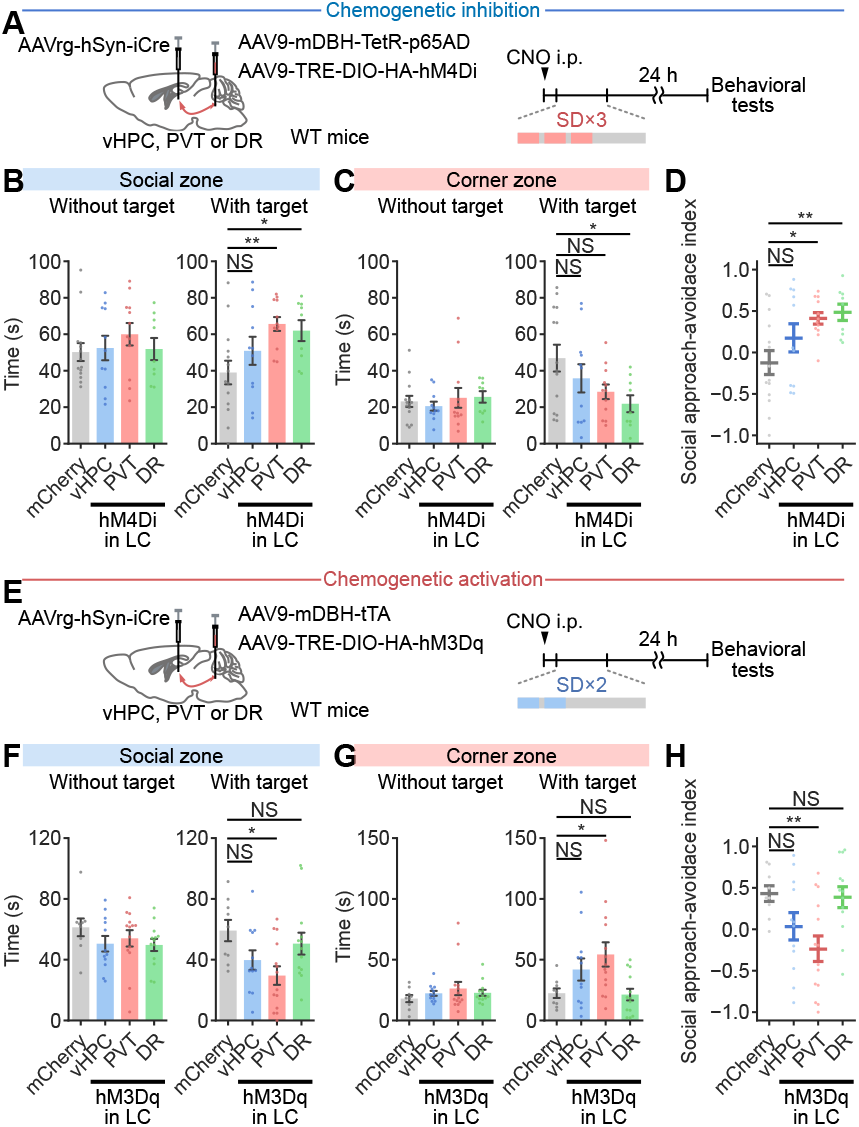
Bidirectional modulation of social avoidance learning mediated by LC^NA^→PVT neurons. Schematic illustrating behavioral experiments for projection target-selective chemogenetic inhibition of LC^NA^ neurons. Mice were subjected to SD×3 episodes 15 min after intraperitoneal administration of 5 mg/kg CNO. The control mice comprised a group of mice expressing mCherry in LC^NA^ neurons retrogradely targeted from the vHPC, PVT, or DR. (**B, C**)Quantification of time spent in the social (B) and corner (C) zones with or without a target mouse. mCherry, n = 14; LC^NA^→vHPC-hM4Di, n = 11; LC^NA^→PVT-hM4Di, n = 12; LC^NA^→DR-hM4Di, n = 11. NS, not significant; *P < 0.05, **P < 0.01, one-way ANOVA followed by Dunnetts test. (**D**) Quantification of the social approach avoidance index. NS, not significant; *P < 0.05, **P < 0.01, one-way ANOVA followed by Dunnetts test. (**E**) Schematic illustrating behavioral experiments for projection target-selective chemogenetic activation of LC^NA^ neurons. Mice were subjected to subthreshold SD×2 episodes 15 min after intraperitoneal administration of 1 mg/kg CNO. The control mice comprised mice expressing mCherry in the pan-LC^NA^ neurons. (**F, G**) Quantification of time spent in the social (F) and corner (G) zones. mCherry, n = 9; LC^NA^→vHPC-hM3Dq, n = 12; LC^NA^→PVT-hM3Dq, n = 14; LC^NA^→DR-hM3Dq, n = 13. NS, not significant; *P < 0.05, one-way ANOVA followed by Dunnetts test. (**H**) Quantification of the social approach avoidance index. NS, not significant; **P < 0.01, one-way ANOVA followed by Dunnetts test. Data are represented as mean *±* SEM.

In contrast, chemogenetic activation yielded distinct outcomes between LC^NA^ subsystems: LC^NA^→PVT neurons during subthreshold SD×2 episodes significantly reduced social investigation behavior and increased the time spent in corner zone, whereas LC^NA^→DR neurons, like LC^NA^→vHPC neurons, failed to elicit any significant behavioral alteration (Figs. 5E–6H, and S8C). These results raise the possibility that noradrenergic inputs to the PVT and adjacent local regions could exert enhancing effects on social avoidance learning. Therefore, we examined the effects of local blockade of beta-adrenergic receptor in the PVT during SD×3 episodes. We found that local infusion of 1 µg of propranolol into the PVT significantly increased the time spent in social interaction zone and the social approach-avoidance index (Figs. S9A–9E). These results suggest that the LC^NA^→PVT subpopulation mediates social trauma-induced avoidance learning, presumably via beta-noradrenergic receptor.

## DISCUSSION

Discrimination of threatening conspecifics is essential for survival and health maintenance, but these processes must be adaptively modulated based on past social experiences to maintain optimal social functioning^45^. In this study, our mesoscopic approaches revealed that traumatic, consecutive social stress engaged a pontothalamic LC^NA^ subpopulation in social avoidance learning, distinct from the subpopulations projecting to hippocampus, amygdala, and other memory storage regions (Figs. 2 and 3). Previous studies have highlighted the central role of the LC^NA^ system in stress-induced anxiety-related behaviors and aversive memory processing. Activation of LC^NA^ neurons by acute stress promotes anxiety-related behaviors via projections to the amygdala^17,25,35^. Moreover, aversive contextual and cued memory processing is modulated by LC^NA^ neurons through outputs to the hippocampus^46^ and amygdala^22^, respectively. Although these neural substrates of general negative emotion have been well characterized, distinct brain networks involved in memory acquisition, arousal regulation, and salience processing can be differentially recruited depending on the type and frequency of stress. Our data demonstrates that the LC^NA^→PVT neurons exhibited response dynamics that tracked the aversive salience of social stimuli (Figs. 4 and S7) and exerted modulatory influences on social trauma-induced avoidance behavior (Fig. 5).

Given that trauma severity including stress uncontrollability is associated with posttraumatic stress disorder^5^, understanding neural responses under different stress conditions is crucial. Our brain-wide activity mapping revealed distinct activation patterns related to stress type (social versus non-social) and stress frequency (single versus consecutive social stress) (Fig. 1). Among all stress paradigms examined, consecutive social stress induced the most widespread activation across the brain. Notably, the LC, a key stress-responsive region^1,34,35^, exhibited heightened activation under consecutive social stress compared to single social stress. This finding suggests that even well-characterized brain regions display distinct activation patterns in response to single versus consecutive social threats. Our dataset of brain-wide responses to various stress paradigms not only underscores these differences but also serves as a valuable resource for future research on stress-responsive neural circuits.

Previous studies have described various projection patterns of LC^NA^ neurons, including targeted^22^, divergent^41^, and combined projections^26,28^, and suggested that distinct LC^NA^ neurons send collateral projections to functionally related target regions^24^. However, the precise organization of their axonal wiring remains unclear. Our projectome data demonstrate that each LC^NA^ subpopulation has highly collateralized yet specific projections to multiple brain regions. The projectome subtypes innervating diencephalic and brainstem structures were largely distinct from another subtype innervating the hippocampal formation and its associated cortical areas (Figs. 3, S5E, S5D, and S6). This comprehensive characterization of collateralization patterns will be crucial for understanding how neuromodulatory systems regulate multiple neural circuits both in parallel and independently.

In addition, we revealed that different projectome subtypes of LC^NA^ neurons have distinct neural coding properties. Our fiber photometry data suggest that LC^NA^→PVT neurons, but not LC^NA^→vHPC neurons, signal dynamic tracking of aversive salience to their downstream regions, including the PVT and DR. Although previous studies have suggested that the LC-PVT circuitry responds to noxious footshock^47^ and modulates arousal^48^, our findings demonstrate an additional aspect of neural processing and functional roles in this pathway. Consistent with our results, previous reports showed similar neural coding properties in the PVT and DR, as both scale their response to aversive predictive cues but not aversive prediction errors^49,50^. Notably, neuronal activity in the PVT is essential for driving associative learning^49,51^. Taken together, our data suggest that the LC^NA^ circuit module connecting to thalamic and brainstem structures plays a critical role in driving aversive social learning by tracking the aversive salience of social stimuli. Future studies will be required to investigate how such response differences emerge between distinct LC^NA^ subpopulations and how dynamic responses in LC^NA^→PVT neurons are shaped, focusing on the afferent connectivity and molecular features of each subpopulation.

In conclusion, our study identified the key neural circuit architecture of the LC^NA^ subsystem that mediates the translation of social trauma into social avoidance behavior. We demonstrated that traumatic social avoidance learning is mediated not by noradrenergic projections to memory-encoding regions such as hippocampus and amygdala, but rather by projections to diencephalic and brainstem regions involved in salience processing. These findings provide critical insights into how traumatic and non-traumatic stress differentially engage distinct functional modules of neuromodulatory circuits, leading to either the impairment or preservation of social functioning.

## Acknowledgements

We are grateful to I. Imayoshi (Kyoto University, Kyoto, Japan) for technical advice on the preparation of 4-OHT solution. This work was supported in part by JSPS KAKENHI Grant Numbers, JP20H03556 (K.S.), JP22K19481 (K.S.), JP23K27331 (A.K.), JP22K19378 (A.K.), JP22H05080 (A.K.), JP23K24205 (T.H.), and JP23K18163 (T.H.); AMED Grant Numbers, JP21wm0525005 (K.S.), JP24wm0625116 (K.S.), JP19gm1310003 (T.Nakazawa), JP21wn0425012 (T.Nakazawa), JP21wm0425010 (T.H.), JP21gm1510006 (T.H.), JP21dm0207117 (H.H.), JP23ama121052 (H.H.), and JP23ama121054 (H.H.); JST, PRESTO Grant Number JP-MJPR23S7 (K.S.); the Collaborative Research Program of Institute for Protein Research, Osaka University, ICR-20-03 (T.H.); and grants from the Takeda Science Foundation (T.Nakazawa, A.K.).

## Author contributions

Conceptualization, K.S.; Methodology, K.S.; Formal Analysis, K.S., S.M., and L.K.; Investigation, K.S., S.M., L.K., Y.F., T.B., H.R., and T.Nitta; Resources, K.S., A.K., T.H., and H.H.; Writing – original draft, K.S.; Writing – review & editing, K.S., T.Nakazawa, A.K., and H.H.; Funding Acquisition, K.S., T.Nakazawa, A.K., T.H., and H.H.; Supervision, K.S.

## MATERIALS AND METHODS

### Animals

All animal care and handling procedures were approved by the Animal Care and Use Committee of Osaka University (approval number, R02-8). All efforts were made to minimize the number of animals used. Male mice aged 7–14 weeks were used for Fos mapping, behavioral, and fiber photometry experiments. For axonal tracing experiments, male and female mice aged 7–14 weeks were used. B6.Cg-Tg(Fos-tTA,Fos-EGFP*)1Mmay/J mice (Fos-EGFP; JAX stock #018306; The Jackson Laboratory, Bar Harbor, ME) and Th^tm1(cre)Te^ mice (TH-Cre; MGI:3056580) were maintained on the C57BL/6J background. Fos^2A-iCreER^ knock-in mice (TRAP2; JAX stock #030323; The Jackson Laboratory) were maintained on the C57BL/6N background. For these transgenic or knock-in mice, heterozygous mice in each mouse line were used for experiments. Wild-type C57BL/6J, C57BL/6N, and retired CD-1 mice were purchased from SLC (Shizuoka, Japan). All behavioral experiments were conducted on the C57BL/J background. Mice were maintained in group housing (3–6 mice per cage) and separated to single-housing at 1 week before the experiments. They were maintained on a 12-h lightdark cycle (lights on at 8:00 a.m.) with controlled room temperature and humidity. Water and food (CMF, Oriental Yeast, Osaka, Japan) were available ad libitum.

### Plasmids

For construction of pAAV-mDBH-tTA plasmid, a promoter region of the mouse DBH gene was cloned from mouse genomic DNA through polymerase chain reaction (PCR), and subsequently it was replaced with mouse oxytocin (OXT) promoter in a pAAV-inverted WPRE (IW)-mOXT-tTA^39^, using basic subcloning techniques. For construction of pAAV-mDBH-TetR-p65AD, DNA fragment coding TetR-p65AD was synthesized and subcloned in a pEX-A2J2 vector by Eurofins genomics (Tokyo, Japan), followed by replacement with the tTA-coding sequence in the pAAV-mDBH-tTA, using basic subcloning techniques.

For AAV transgene plasmids carrying Cre-dependent expression cassettes of pal-mScarlet, jGCaMP7s, and DREADDs, coding sequences of these genes were inserted into a linearized pAAV-TRE-DIO (obtained by restriction enzyme digestion of pAAV-TRE-DIO-FLPo^52^; a gift from Minmin Luo; Addgene plasmid # 118027; http://n2t.net/addgene:118027; RRID:Addgene_118027) using basic plasmid ligation or the In-Fusion cloning system (639650, Clontech/Takara). These coding sequences were cloned by PCR from pAAV-TRE-pal-mScarlet^39^, pAAV-hSyn-jGCaMP7s^43^ (a gift from Douglas Kim & GENIE Project; Addgene plas-mid #104487; http://n2t.net/addgene:104487; RRID:Addgene_104487), pAAV-hSyn-DIO-HA-hM4Di-IRES-mCitrine (a gift from Bryan Roth; Addgene plasmid #50455; http://n2t.net/addgene:50455; RRID:Addgene_50455), pAAV-hSyn-DIO-HA-hM3Dq-IRES-mCitrine (a gift from Bryan Roth; Addgene plasmid #50454; http://n2t.net/addgene:50454; RRID:Addgene_50454), and pAAV-EF1a-DIO-EYFP (a gift from Karl Deisseroth; Addgene plasmid #20296; http://n2t.net/addgene:20296; RRID:Addgene_20296).

For AAV transgene plasmids carrying a Cre expression cassette, pAAV-hSyn-iCre-WPRE was first prepared by inserting a PCR-cloned DNA fragment coding iCre from pCAG-iCre^53^ (a gift from Wilson Wong; Addgene plasmid #89573; http://n2t.net/addgene:89573; RRID:Addgene_89573) into a linearized pAAV-hSyn using In-Fusion cloning system. The WPRE sequence in the pAAV-hSyn-iCre-WPRE was removed by digestion with ClaI followed by self-ligation. The linearized pAAV-hSyn was obtained by restriction enzyme digestion of pAAV-hSyn-DIO-mCherry (a gift from Bryan Roth; Addgene plasmid #50459; http://n2t.net/addgene:50459; RRID:Addgene_50459).

### Viral vector production

Lenti-X 293T cells (632180, Clontech, San Jose, CA) were used to produce the AAV. Cells were cultured in Dulbeccos modified Eagles medium with high glucose and GlutaMAX supplement (DMEM, high glucose, GlutaMAX™ Supplement; 10566024, Thermo Fisher Scientific, Waltham, MA) containing 10% fetal bovine serum (Sigma-Aldrich, St Louis, MO) at 37°C with 5% CO_2_. AAV vectors were produced using the helper-free triple transfection procedure, as previously described^39,54^. We used AAV serotype 9 and 2-retro capsids carried on the replication and capsid proteins plasmids, a pAAV2/9n plasmid (a gift from James M. Wilson; Addgene plasmid # 112865; http://n2t.net/addgene:112865; RRID:Addgene_112865) and a rAAV2-retro helper plasmid^42^ (a gift from Alla Karpova & David Schaffer; Addgene plasmid # 81070; http://n2t.net/addgene:81070; RRID:Addgen_81070), respectively. An AAV transgene plasmid, a replication and capsid protein plasmid, and a pHelper plasmid (VPK-400-DJ, Cell Biolabs, San Diego, CA), which supplied the adenovirus gene products required for AAV production, were cotransfected into Lenti-X 293T cells (632180, Clontech, San Jose, CA) using polyethyleneimine (PEI MAX; 24765, Polyscience Inc., Warrington, PA). Cells were cultured in Dulbeccos modified Eagles medium with high glucose and GlutaMAX supplement (DMEM, high glucose, GlutaMAX Supplement; 10566024, Thermo Fisher Scientific, Waltham, MA) containing 10% fetal bovine serum (Sigma-Aldrich, St Louis, MO) at 37°C with 5% CO_2_. The AAV vectors were collected from the cells and media by ultracentrifugation using an iodixanol step gradient (15%, 25%, 40%, and 60%; Optiprep, 1893, Cosmo Bio, Tokyo, Japan) and were replaced and concentrated with phosphate-buffered saline (PBS) containing 0.001% (v/v) Pluronic F-68 (24040032, Thermo Fisher Scientific) via ultrafiltration using an Amicon Ultra-15 centrifugal filter (100-kDa cutoff; UFC910024, Millipore). The viral titers were quantified by a quantitative real-time PCR using GoTaq qPCR Master Mix (Cat# A6001, Promega, Madison, WI), with a linearized AAV genome plasmid serving as a standard.

### Stereotaxic surgery

Mice were deeply anesthetized via an intraperitoneal injection of a mixture containing 0.75 mg/kg medetomidine (Nippon Zenyaku Kogyo, Fukushima, Japan), 4 mg/kg midazolam (Sandoz Pharma, Basel, Switzerland), and 5 mg/kg butorphanol (Meiji Seika Pharma, Tokyo, Japan). Next, they were placed in a stereotaxic instrument (RWD Life Science Co., LTD, Guangdong, China) with a 37°C heat pad. Viral injections were performed using a Neuros Syringe with a 33-gauge needle (65460-06, Hamilton, Reno, NV) or a gastight syringe (1701RN, Hamilton) with a glass needle compression fitting (55750-01, Hamilton) and a self-made grass micropipette, which was prepared using a glass capillary (GD-1, Narishige, Tokyo, Japan) and a glass capillary puller (PC-100, Narishige). Injection volumes and speeds were controlled using an UltraMicroPomp3 with SMARTouch Controller (World Precision Instruments, Sarasota, FL). The needle was left in the target brain region for 3 min before withdrawal and slowly removed to avoid backflow. After stereotaxic surgery, the mice immediately received an intraperitoneal injection of atipamezole (7.5 mg/kg; Nippon Zenyaku Kogyo) and gentamicin (10 mg/kg; Sigma-Aldrich). One day after the surgery, the mice received an intraperitoneal injection of buprenorphine (0.1 mg/kg; Otsuka Pharma, Tokyo, Japan) for pain relief.

### Surgery for labeling cell bodies or axons

C57BL/6 mice and TRAP2 mice aged 7–9 weeks on the day of stereotaxic surgery were used. For labeling pan-LC^NA^ neurons, AAV9-mDBH-tTA at 3–5 ×10^12^ viral genomes (vg)/ml were mixed with AAV9-TRE-mNeonGreen (prepared from pAAV-TRE-mNeonGreen^55^; a gift from Viviana Gradinaru; Addgene plasmid #99131; http://n2t.net/addgene:99131; RRID:Addgene_99131) or AAV9-TRE-pal-mScarlet at 8–10×10^12^ vg/ml in a 1:1 volume ratio. The viral cocktail at a volume of 500 nl per side was injected into the LC at anteroposterior (AP) −5.5 mm, mediolateral (ML) −0.95 mm, and dorsoventral (DV) −4.25 mm from the bregma using a 33 gauge 12°-beveled needle and a Neuros Syringe (65460-06, Hamilton, Reno, NV). At two weeks after surgery, brain tissues were collected as described in the section Tissue preparation. For neuronal activation-dependent labeling, AAV9-mDBH-tTA (3–5×10^12^ vg/ml) and AAV9-TRE-DIO-pal-mScarlet (1×10^13^ vg/ml) were mixed at a 1:1 volume ration, and 500 nl of the viral cocktail was injected into the LC of TRAP2 mice. For projection target-selective labeling, 500 nl of the viral cocktail of AAV9-mDBH-tTA and AAV9-TRE-DIO-pal-mScarlet was injected into the LC, and 200 nl of AAVrg-hSyn-iCre (4×10^12^ vg/ml) was injected into the vHPC (AP −3.4 mm, ML −3.0 mm, DV −4.2 mm from the bregma), PVT (AP −1.6 mm, ML ± 0.0 mm, DV −3.0 mm from the bregma), or DR (AP −4.3 mm, ML ±0.0 mm, DV −3.0 mm from the bregma). For the PVT and DR, stereotaxic injections to the target coordinates were performed at a 20°angle to the vertical axis in order to avoid damaging the superior sagittal sinus. At three weeks after surgery, brain tissues were collected as described in the section Tissue preparation.

### Surgery for DREADDs expression

C57BL/6J mice and TH-Cre mice aged 7–8 weeks on the day of stereotaxic surgery were used. For hM4Di expression in TH-Cre mice, 500 nl of AAV9-hSyn-DIO-hM4Di-mCherry (2×10^13^ vg/ml) or AAV9-hSyn-DIO-mCherry (1×10^13^ vg/ml) was bilaterally injected into the LC (AP −5.5 mm, ML ±0.95 mm, DV −4.25 mm from the bregma) using a 33 gauge 12°-beveled needle. For hM4Di expression in pan-LC^NA^ neurons in wild-type C57BL/6 mice, AAV9-mDBH-TetR-p65AD (5×10^12^ vg/ml) and AAV9-TRE-HA-hM4Di (7×10^12^ vg/ml) were mixed at a 1:1 volume ratio, and 500 nl of the viral cocktail was bilaterally injected into the LC. For projection target-selective chemogenetic inhibition and activation, total 400 nl of AAVrg-hSyn-iCre (4×10^12^ vg/ml) was injected into the vHPC (AP −3.4 mm, ± ML 3.0 mm, DV −4.2 mm from the bregma), PVT (AP −1.6 mm, ML ±0.0 mm, DV −3.0 mm from the bregma), or DR (AP −4.3 mm, ML ±0.0 mm, DV −3.0 mm from the bregma). For projection target-selective hM4Di expression, AAV9-mDBH-TetR-p65AD (5×10^12^ vg/ml) and either AAV9-TRE-DIO-mCherry (1×10^13^ vg/ml) or AAV9-TRE-DIO-HA-hM4Di (5×10^12^ vg/ml) were mixed at a 1:1 volume ratio, and 500 nl of the viral cocktail was bilaterally injected into the LC. For projection target-selective hM3Dq expression, 500 nl of the viral cocktail of AAV9-mDBH-tTA and either AAV9-TRE-mCherry (5–9 10^12^ vg/ml) or AAV9-TRE-DIO-HA-hM3Dq-P2A-EYFP (9–10 10^12^ vg/ml) was bilaterally injected into the LC.

### Surgery for fiber photometry

C57BL/6J mice aged 7–8 weeks received stereotaxic injections of a cocktail of AAV9-mDBH-tTA and AAV9-TRE-DIO-jGCaMP7s (7 ×10^12^ vg/ml) at a mixed volume ratio of 1:1 into the unilateral LC as well as 200 nl of AAVrg-hSyn-iCre into the ipsilateral vHPC, PVT, or DR. Subsequently, a fiber-optic cannula (400-µm diameter, NA 0.37, fiber length 4.0 mm; MFC_400/470-0.37_4.0mm_MF1.25_FLT, Doric Lenses, Québec, Canada) was implanted over the right LC (AP −5.5 mm, ML ± 0.90 mm, DV −3.5 mm from the bregma). The ferrule of the fiber-optic cannula was attached to the skull surface using Super-Bond (Sun Medical Co. Ltd., Shiga, Japan), and was further anchored with an anchor screw (AN-3, Eicom, Kyoto, Japan) and dental cement (UNIFAST III, GC Corp., Tokyo Japan). The top of the fiber-optic cannula was covered with a low-toxicity silicon adhesive (KWIK-SIL, World Precision Instruments) to keep it clean.

### Surgery for drug infusion

For local drug infusion into the PVT, a guide cannula (0.3-mm outer diameter, 0.4-mm inner diameter, 3.5-mm length; AG-3,5(T), Eicom) with a dummy cannula (AD-3.5(T), Eicom) was implanted targeting the PVT (AP −1.6 mm, ML ± 0.0 mm, DV −3.0 mm from the bregma) at a 10°angle to the vertical axis to avoid damaging the superior sagittal sinus. The cannula was attached to the skull surface using Super-Bond and further anchored with an anchor screw and dental cement.

### Stress models

#### Social defeat stress for behavioral tests

Aggressive CD-1 mice were pre-screened for aggressive behavior as previously described^15,56^ with minor modifications. We extended the screening period to 5 days, and aggressive behavior of CD-1 was evaluated during the last 3 days. CD-1 mice which met the criteria (attack latencies < 60 s, and two or more attacks during a 3-min session on at least 2 consecutive days) were used as aggressor mice. All behavioral tests were performed on the C56BL/6J background, and experimental mice were singly housed at 1 week before social defeat stress. Experimental mice were placed into the aggressors home cage for 10 min per social defeat stress episode. For repeated social defeat episodes, each experimental mouse was exposed to a different, unfamiliar aggressor for each separate encounter. For SD×5, 10-min social defeat stress episodes were repeated five times with 2-min intervals. For S×D 2, a clear perforated divider was inserted between the experimental and aggressor mice 3 min after the onset of each social defeat stress episode, which remained there for the rest of the episode. For SD×3, the divider was inserted 5 min after the onset of each social defeat stress episode and remained there for the remaining 5 min. In case of any signs of excessive wounding, the stress session was stopped. For chemogenetic inhibition experiments using hM4Di, 5 mg/kg CNO was intraperitoneally administrated 15 min before the onset of SD×3. For systematic inhibition of the adrenergic receptors, 1 mg/kg prazosin (P7791, Sigma-Aldrich), 10 mg/kg propranolol (P0884, Sigma-Aldrich), or saline were intraperitoneally administrated 30 min before the onset of SD×3. For chemogenetic activation experiments using hM3Dq, 1 mg/kg CNO was intraperitoneally administrated 15 min before the onset of SD×2. For local drug infusion into the PVT, 100 nl of 10 mg/ml propranolol (total 1 µg per mouse) or phosphate-buffered saline (PBS; 045-29795, Fujifilm-Wako, Osaka, Japan) was infused via the implanted cannula at 100 nl/min at 15 min before the onset of SD×3. After social interaction tests, 150 nl of Hoechst 33342 (10 µg/ml) was infused via the implanted cannula to visualize the infusion site.

#### Stress models for Fos expression analysis

Fos-EGFP mice aged 8–10 weeks at the start of single housing were used, and single housing of each experimental mouse was started 7 days before the fixation. To habituate mice to the experimental environment and avoid environment-induced Fos expression, the mice were placed in the experimental room for 1 h daily for 3 consecutive days before stress exposure and fixation. The last habituation session was finished at least 24 h prior to the stress exposure. For SD×1 and SD×5 conditions, mice were subjected to these stress as described in the subsection entitled Social defeat stress for behavioral tests. For naïve home cage condition, mice were transferred to the experimental room and stayed in their home cage there for 1 h. For novel environment conditions, mice were placed in the novel cage without CD1 mouse for 1 h. For restraint stress conditions, mice were placed in a 50-ml polypropylene tube with holes at the tips and lateral sides for unrestricted breathing for 1h. For footshock conditions, mice were placed in a fear conditioning chamber with an electrical grid floor (Med Associates, Fairfax, VT,) for 45 min and received 10 footshocks (0.3 mA or 0.8 mA, 0.5 s) paired with pure tones (CS+, 15,000 Hz, 20 s). Footshocks were delivered after the offset of CS+. The other tone (CS−, 7,500 Hz, 20 s) was presented intermittently, following the paring of CS+ with the footshock. The interval between footshock presentations was randomized between 2–6 min. At the end of either condition, mice were returned to their housing room and kept there for an additional 2 h for EGFP expression^57^. For Fos expression analysis in TH+ cells in the LC, C57BL/6J mice aged 8 weeks were subjected to SD×5 and slightly modified SD×1 conditions. For modified SD×1, mice were exposed to 10 min of social defeat stress followed by 50 min of sensory contact with the same aggressor using a clear perforated divider. Mice were fixed 2 h after the onset of social defeat stress. Finally, brain tissues were collected as described in the Tissue preparation section below.

#### Social defeat stress for fiber photometry

On the test day, mice were habituated in the experimental room for at least 30 min before fiber photometry recording. A single stress session comprised three phases: a baseline phase in an aggressors home cage without the aggressor (2 min), a sensory contact phase with the aggressor mouse placed in a cage (5 min), and a physical contact phase with the freely behaving aggressor mouse (3 min). During the physical contact phases, in order to balance the number of attacks received across mice, we limited the maximum number of continuous attacks lasting more than 5 s to five per mouse in each session by inserting a clear perforated divider into the cage. The stress session was repeated five times with 2-min intervals.

### Behavioral tests

#### Social interaction test

The social interaction test was performed as previously described^15,56^. Under red-light conditions, which were maintained throughout the experiment, the experimental mouse was habituated to the test room in its home cage for at least 30 min. The test comprised two 150-s sessions with or without a target CD1 male mouse. For the first session, the experimental mouse was placed on the open field (42 cm width×42 cm depth×40 cm height; Brain Science Idea, Osaka, Japan) containing an empty wire-mesh cage (10 cm×6.5 cm; Brain Science Idea) that was located at one end of the field. For the second session, the experimental mouse was placed in the same field as in the first session but with an unfamiliar target CD1 mouse in the wire-mesh cage. Between these two sessions, the experimental mouse was returned to its home cage for approximately 1 min. The open field and wire mesh cage were cleaned with 70% ethanol after each test. The interaction zone was defined as a 14 cm ×24 cm area around the wire-mesh cage, and the corner zones were defined as two 9 cm×9 cm areas, which were each located in one of the two corners opposite to the wire-mesh cage. The tests were recorded using a video camera (HC-VX992M, Panasonic, Osaka, Japan), and the time spent in each zone was analyzed using ANY-maze video-tracking software (version 4.99 m, Stoelting, Wood Dale, IL). To calculate the social approach-avoidance index, difference between times spent in the interaction and corner zones (interaction zone corner zone) was divided by sum of times spent in these zones (interaction zone + corner zone) in the second session with a target CD1 mouse.

#### Open field test

The open field test was performed in a square box (42 cm width ×42 cm depth ×40 cm height), as previously described^15^, with minor modifications. Under 40–60 lux light intensity conditions, which were maintained throughout the experiment, the experimental mouse in its home cage was habituated to the test room for at least 30 min. Each experimental mouse was placed in one corner of the open field and allowed to move freely for 5 min. The open field was cleaned with 70% ethanol after each test. The tests were recorded using a video camera, and the total distance traveled and time spent in the center zone (21 cm ×21 cm) were analyzed using ANY-maze video-tracking software.

#### Elevated plus maze test

The elevated plus maze test was performed as previously described^15^, with minor modifications. Under 40–60 lux light intensity conditions, which were maintained throughout the experiment, the experimental mouse in its home cage was habituated to the test room for at least 30 min. The elevated plus maze apparatus (Brain Science Idea), which comprises two open arms (25 cm length × 8 cm width) and two closed arms (25 cm length×8 cm width, surrounded by an opaque wall with a height of ×20 cm) placed at a height of 50 cm from the ground, was used. Each experimental mouse was placed at the end of one closed arm and allowed to move freely for 5 min. The apparatus was cleaned with 70% ethanol after each test. The tests were recorded using a video camera, and the time spent in the open arms and the number of entries into the open arms were analyzed using ANY-maze video-tracking software. Mice that fell from the maze during the test were excluded from the analysis.

### Neuronal activation-dependent labeling of axons

To induce CreER^T2^-mediated recombination, 4-OHT (H6278, Sigma-Aldrich) was first dissolved in dimethyl sulfoxide (DMSO; D2650, Sigma-Aldrich) and further diluted into PBS containing 1% Tween 80 (Polyoxyethylene Sorbitan Monooleate; 35703-62, Nacalai Tesque) as previously described^58^. The final concentration of 4-OHT solution was 2.5 mg/ml 4-OHT, 5% DMSO, and 1% Tween 80 in PBS. At 10 days after surgery, TRAP2 mice intraperitoneally received 25 mg/kg 4-OHT at the end of the stress episodes. Three weeks after 4-OHT administration, brain tissues were collected as described in the section Tissue preparation.

### Tissue preparation

Mice were deeply anesthetized by intraperitoneal injection of an anesthetic cocktail containing 0.75 mg/kg medetomidine, 4 mg/kg midazolam, and 5 mg/kg butorphanol. Anesthetized mice were transcardially perfused with 10 ml of saline followed by 10 ml of 4% paraformaldehyde (PFA) (Nacalai Tesque, Kyoto, Japan) dissolved in PBS (137 mM NaCl, 8.1 mM Na_2_HPO_4_/12H_2_O, 2.7 mM KCl, 1.47 mM KH_2_PO_4_; pH 7.4). For whole-brain imaging experiments, brain tissues were excised and immersed in 4% PFA dissolved in PBS for at least 2 days and susequently transferred and stored in 0.05% sodium azide (S2002, Sigma-Aldrich) dissolved in PBS at 4°C until use. For immunohistochemical analysis, the fixed brains were cryoprotected in 20% (w/v) sucrose dissolved in PBS for 2 days, embedded in OCT compound (Sakura Finetek, Tokyo, Japan), and quickly frozen on liquid nitrogen. The frozen brain tissues were stored at −30°C until use.

### Whole-brain imaging

Whole-brain imaging was performed as previously described^30,31^ using a serial-section imaging system comprising a vibrating microtome and a spinning disk confocal microscope, which was named FAST. Each brain tissue embedded in 4% agarose gel dissolved in 0.1 M phosphate buffer, pH 7.4, was placed in a sample chamber filled with PBS. Serial coronal section images were acquired at a lateral resolution of 1 µm/pixel and an axial resolution of 5 µm/pixel, by repeating confocal optical sectioning imaging and mechanical sectioning with a vibrating microtome at 50-µm thickness. All whole-brain images were acquired using two color channels: fluorescent protein and tissue autofluorescence signals. For Fos-EGFP mouse brains, EGFP signals were collected using excitation with a 488 nm laser and detection with a 520/35 nm emission filter, and tissue autofluorescence signals were collected using excitation with a 488 nm laser and detection with a 617/73 nm emission filter. For mScarlet-labeled mouse brains, mScarlet signals were collected using excitation with a 561 nm laser and detection with a 617/73 nm emission filter, and tissue autofluorescence signals were collected using excitation with a 488 nm laser and detection with a 520/35 nm emission filter.

### Whole-brain data analysis

#### Mapping Fos-EGFP^+^ cells

Serial section images acquired using FAST at a resolution of 1×1×5 µm were processed and analyzed using custom python codes. The python codes will be publicly available. This pipeline was implemented using scikit-image^59^ (https://scikit-image.org/), OpenCV^60^ (https://opencv.org/; opencv-python), Scipy^61^ (https://scipy.org/), and Numpy^62^ (https://numpy.org/). Section images for tissue autofluorescence and Fos-EGFP images were separately reconstructed, and subsequently Fos-EGFP signals were extracted by subtracting tissue autofluorescence signals. Extracted EGFP images were further subjected to boundary separation of adjacent cells by calculating the difference of Gaussian, binarization, signal size filtering, and water-shedding. Next, the centroids of each binarized cell were detected and quantified in 50-µm cubic increments across the whole brain. Subsequently, these quantification results were then stored as pixel intensities in a downscaled multi-page TIFF file with a 50-µm isotropic resolution and 16-bit depth.

#### Mapping axons

Serial section images at a resolution of 1 ×1 ×5 µm were processed and analyzed using custom python codes. Section images for autofluorescence and mScarlet-labeled axon images were separately reconstructed. To remove small granular autofluorescence signals, which are commonly observed in both autofluorescence and mScarlet channels, each channel was initially subjected to background removal by calculating the difference of Gaussian, and the small granular autofluorescence signals were removed by subtracting the intensity-adjusted autofluorescence channel from the mScarlet channel. Extracted axon images were further subjected to skeletonization of signals, and pixel counts of skeletonized axons in mScarlet-labeled axon images were quantified in 50-µm cubic increments across the whole brain. Similar to Fos-EGFP mapping analysis, quantification results were subsequently stored as pixel intensities in a downscaled multi-page TIFF file with a 50-µm isotropic resolution and 16-bit depth.

#### Registration

The autofluorescence images were used for registration to the Allen reference brain CCFv3^32^ space. To correct optical vignetting in each field of view (FOV), autofluorescence images were subjected to flat-field correction and reduction of low-frequency content by dividing the median-filtered image by the image filtered with a large Gaussian kernel (implemented in OpenCV). The resultant FOVs were stitched, and the coronal section images were reconstructed and rescaled to a 50-µm isotropic resolution. Additionally, the reference atlas images were processed to reduce low-frequency content using custom python codes. Registration was performed using 3D Slicer^63^ and elastix^64^ extension. For anatomical region-based analysis, the atlas annotation data were registered to the autofluorescence tissue images, and the total cells, axon pixel counts, and volumes of each annotated region were quantified. The total value of the left and right brain regions for Fos-EGFP mice were calculated. For axon-labeled brains, the contralateral cerebral hemisphere of the virus injection side was excluded due to the sparse contralateral innervations. Principal component analysis (PCA; implemented in Scikit-learn; https://scikit-learn.org/stable/) was performed on the log-transformed Fos-EGFP+ cell densities in each brain region. Brain regions were classified based on axon densities using k-means clustering (implemented in Scikit-learn). For voxel-based analysis, maps of P-values for between-group comparisons were generated using a t-test assuming unequal variances (implemented in SciPy), and subsequently, maps for differences between groups were shown with diverging colormap.

#### Fiber photometry

Fiber photometry recordings were performed using a custom-built fiber photometry system assembled as previously described^19,65,66^ with minor modifications. Fiber-coupled 405 nm and 470 nm LEDs (405 nm, M405F3; 470 nm, M470F4, Thorlabs, Newton, NJ) were used for the GCaMP isosbestic and fluorescence excitations, respectively. Excitation lights were focused by a 20 ×/NA 0.4 objective lens (PLN20X, Olympus, Tokyo, Japan) to the tip of the 400-µm core, NA 0.37 fiber patch cord (MFP_400/440/1100-0.37_2m_FCM-MF1.25_LAF, Doric lens). The other end of the fiber patch cord was connected to the implanted optical fiber cannula via an interconnect (ADAL3, Thorlabs). The fluorescence signals were separated from excitation lights usig a long-pass dichroic mirror (DMLP505R, Thorlabs) and focused at a CMOS camera (CS235MU, Thorlabs) by an achromatic lens (focal length = 35 mm; AC254-035-A-ML, Thorlabs). The camera was regulated by the ThorCam software and its python SDK (Thorlabs). The camera acquired images at 60 Hz with switching alternately between 405-nm and 470-nm excitation, which yielded a 30 Hz sampling rate for each fluorescence signal.

Data analyses of fiber photometry recordings were performed using a python code, photometry preprocessing (https://github.com/ThomasAkam/photometry_preprocessing), with minor modifications. Regions of interest for the crosssection of the optical fiber were manually defined on the acquired images, and the mean fluorescence intensity was calculated per image. To calculate temporal changes in fluorescence intensity, ΔF/F_0_ (ΔF, fluorescence intensity with subtracted isosbestic signal intensity; F_0_, baseline fluorescence intensity) was obtained as follows. The baseline signal intensities for the isosbestic point and fluorescence signals were estimated by low-pass filtering and subtracted from each signal. To correct for motion artifacts, the isosbestic signals were scaled to the fluorescence signals by lasso regression (implemented in Scikit-learn) and subtracted from the fluorescence signals, which resulted in the output of ΔF values. Then, the ΔF values were divided by the low-pass filtered baseline signals in the fluorescence (F_0_) values. The ΔF/F_0_ values in each session were normalized through z-score-normalization with the mean and standard deviation of the values during the habituation phase (first 2 min). Mean peri-event activities within 3 s before and after the onset of behavioral events were quantified.

#### Immunohistochemistry

Tissue sections were prepared at 20- or 30-µm-thickness using cryostat (CM1860, Leica Biosystems, Nussloch, Germany). Next, they were subjected to flee-floating staining or mounted on glass slides (MAS-04, Matsunami Glass Ind., Ltd, Osaka, Japan) and stained on the slides as described below. After immunostaining, the sections on the glass slides were coverslipped with ProLong Glass antifade mountant (P36980, Thermo Fisher Scientific).

For Fos expression analysis in TH+ neurons, sections were First, 20-µm-thick sections including the LC were mounted on glass slides (eight sections per slide), followed by immunostaining. Sections were incubated with blocking buffer consisting of 1% bovine serum albumin (01860-07, Nacalai Tesque) dissolved in PBS containing 0.5% Triton X-100 (Polyoxyethylene(10) Octylphenyl Ether; 168-11805, Fujifilm-Wako) for 1 h at room temperature for blocking and permeabilization. After blocking and permeabilization, sections were incubated with rabbit monoclonal anti-Fos antibody (1:1,000 dilution; #2250, Cell Signaling Technology, Danvers, MA) and chicken polyclonal anti-TH antibody (1:1,000 dilution; ab76442, abcam) at 4°C overnight for the primary antibody reaction. For the secondary antibody reaction, sections were incubated with Alexa 488-conjugated goat anti-chicken immunoglobulin Y (IgY) and Alexa 568-conjugated goat anti-rabbit immunoglobulin G (IgG) antibodies (1:1,000 dilution; ab150169 and ab175471, respectively, abcam) diluted in buffer containing 1 µg/ml Hoechst 33342 at final concentration.

For immunohistochemical verification of the cell type-specific expression of mNeonGreen in LC^NA^ neurons, sections were first collected into PBS, followed by free-floating staining. Sections were incubated with 1% bovine serum albumin (01860-07, Nacalai Tesque) dissolved in PBS containing 0.5% Triton X-100 (Polyoxyethylene(10) Octylphenyl Ether; 168-11805, Fujifilm-Wako) for 1 h at room temperature for blocking and permeabilization. For the primary antibody reaction, sections were incubated with rabbit polyclonal anti-TH antibody (1:1,000 dilution; AB152, Merck Millipore) diluted in blocking buffer at 4°C overnight. For the secondary antibody reaction, the sections were incubated with Alexa 568-conjugated goat anti-rabbit IgG antibody (1:1,000 dilution; ab175471, abcam, Cambridge, UK) diluted in blocking buffer containing 1 µg/ml Hoechst 33342 (346-07951, Fujifilm-Wako) at final concentration for nuclear staining.

For fiber photometry experiments, validation of the optic-fiber position and jGCaMP7s expression in LC^NA^ neurons were performed. First, 30-µm-thick sections including the LC were mounted on glass slides (eight sections per slide), followed by immunostaining. The same buffers with free-floating staining were used. After blocking and permeabilization, sections were incubated with rabbit polyclonal anti-GFP antibody (1:1,000 dilution; 598, MBL, Tokyo, Japan) and chicken polyclonal anti-TH antibody (1:1,000 dilution; ab76442, abcam) at 4°C overnight for the primary antibody reaction. For the secondary antibody reaction, sections were incubated with Alexa 488-conjugated goat anti-rabbit IgG and Alexa 568-conjugated goat anti-chicken IgY antibodies (1:1,000 dilution; ab150077 and ab175477, respectively, abcam) diluted in buffer containing 1 µg/ml Hoechst 33342 at final concentration.

For DREADDs experiments, validation of DREADDs expression in LC^NA^ neurons was performed. Sections were collected into PBS and subjected to free-floating staining. For the primary antibody reaction, sections were incubated with rabbit monoclonal anti-HA tag antibody (1:1,000 dilution; 3724, Cell Signaling Techynology, Danvers, MA) and chicken polyclonal anti-TH antibody (1:1,000 dilution) at 4°C overnight. For the secondary antibody reaction, sections were incubated with Alexa 568-conjugated goat anti-rabbit IgG and either Alexa 488-conjugated goat anti-chicken IgY (in the case of HA-hM4Di expression) or Alexa 647-conjugated goat anti-chicken IgY (in the case of HA-hM3Dq-P2A-EYFP expression) antibodies (1:1,000 dilution; ab150169 and ab150171, respectively, abcam) in buffer containing 1 µg/ml Hoechst 33342 at final concentration.

#### Quantification and statistical analysis

All data are presented as mean ±SEM, except for box plots. Statistical analyses were conducted on Python 3.9 or later, using Scipy and scikit-posthocs (https://scikit-posthocs.readthedocs.io/en/latest/index.html) libraries. Outliers in behavioral tests were detected by Grubbs test and removed from statistical analyses. Students t-test, paired t-test, one-way ANOVA, twoway repeated measures ANOVA, Dunnetts test, Tukeys test, HolmSidak correction, Bonferroni correction, false discovery rate (FDR) correction, KruskalWallis test, and Steel-Dwass test were applied, as appropriate. All statistical tests were performed as two-tailed. Statistical significance was set at a P-value < 0.05 (*P < 0.05, **P < 0.01, ***P < 0.001).

## Supplementary figures

**Fig. S1.**
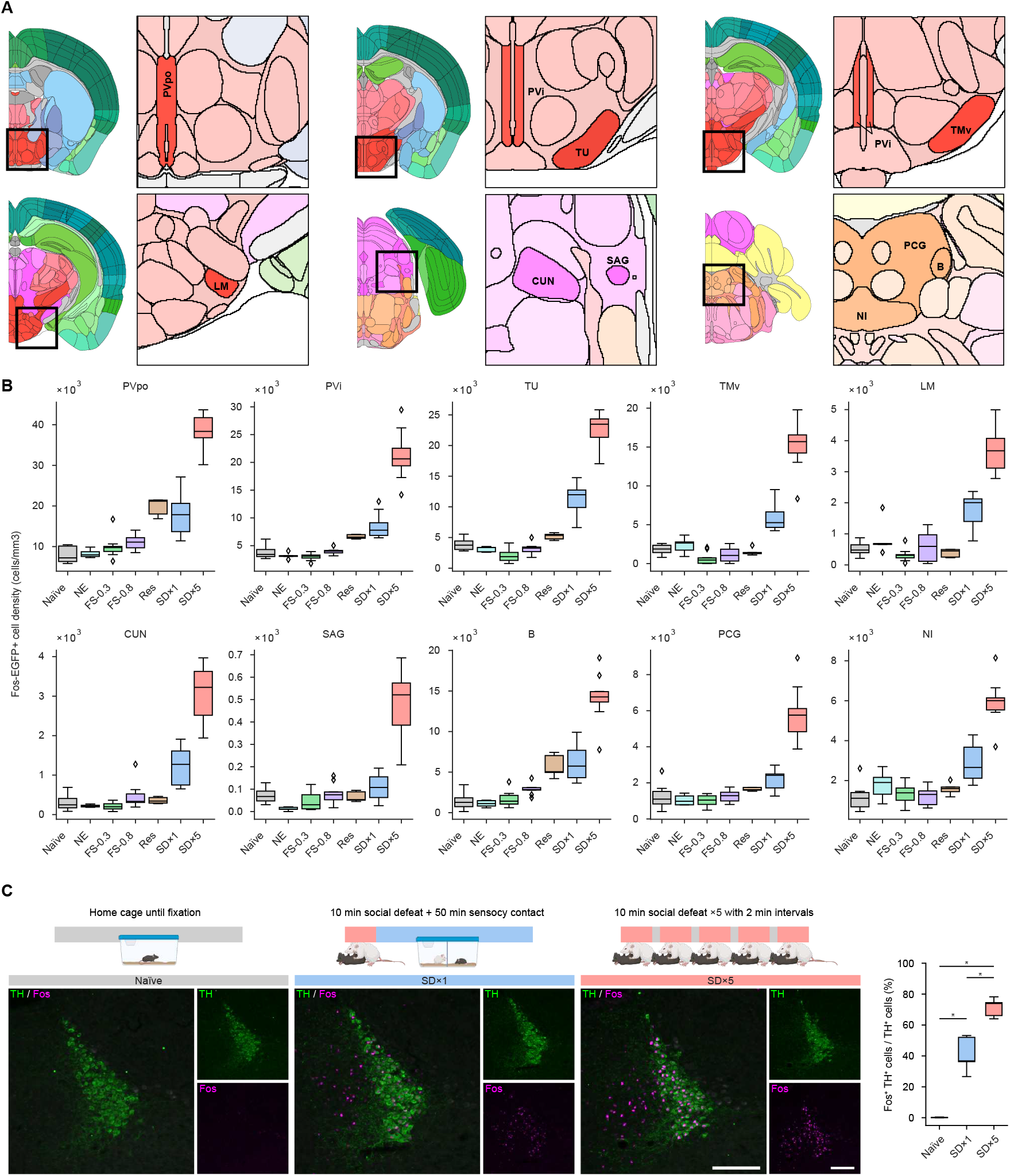
Distinct activity changes in brain regions by different stress conditions. (A) Coronal atlas views of brain regions showing statistical differences in Fos-EGFP+ cells between SD×1 and SD×5: higher effect size (Cohens d > 3), fold change (SD×5 over SD×1) > 2, and adjusted p-value < 0.05. Allen brain atlas CCFv3 was used to classify brain regions. PVpo, periventricular hypothalamic nucleus; PVi, periventricular hypothalamic nucleus, intermediate part; TU, tuberal nucleus; TMv, tuberomammillary nucleus, ventral part; LM, lateral mammillary nucleus; CUN, cuneiform nucleus; SAG, nucleus sagulum; PCG, pontine central gray; B, barringtons nucleus; NI, nucleus incertus. (**B**) Quantification of Fos-EGFP density in brain regions indicated in (a) following exposure to different stress types and magnitudes. Naïve, n = 10; SD×1, n = 10; SD×5, n = 10; NE, n = 5; Res, n = 5, FS-0.3, n = 10; FS-0.8, n = 10. (**C**) Representative images of immunohistochemical analysis and quantification of stress-induced Fos expression in LC^NA^ neurons. For SD×1, mice were subjected to 10 min social defeat stress followed by 50 min sensory contact separated with a clear perforated divider. The sections were immunostained for TH (green) and Fos (magenta). The percentage of Fos+ cells among TH+ cells was quantified. Naïve, n = 5; SD×1, n = 5; SD×5, n = 5. *P < 0.05; Kruskal-Wallis test followed by Steel-Dwass test. For boxplots, boxes indicate the interquartile range with median lines, whiskers extend to 1.5 times the interquartile range, and points represent outliers beyond these ranges. Scale bar, 200 µm. The illustrations of the coronal brain sections in (A) were adapted from the Allen Brain Reference Atlas (https://atlas.brain-map.org). The illustrations of mice in (C) were created with BioRender.

**Fig. S2.**
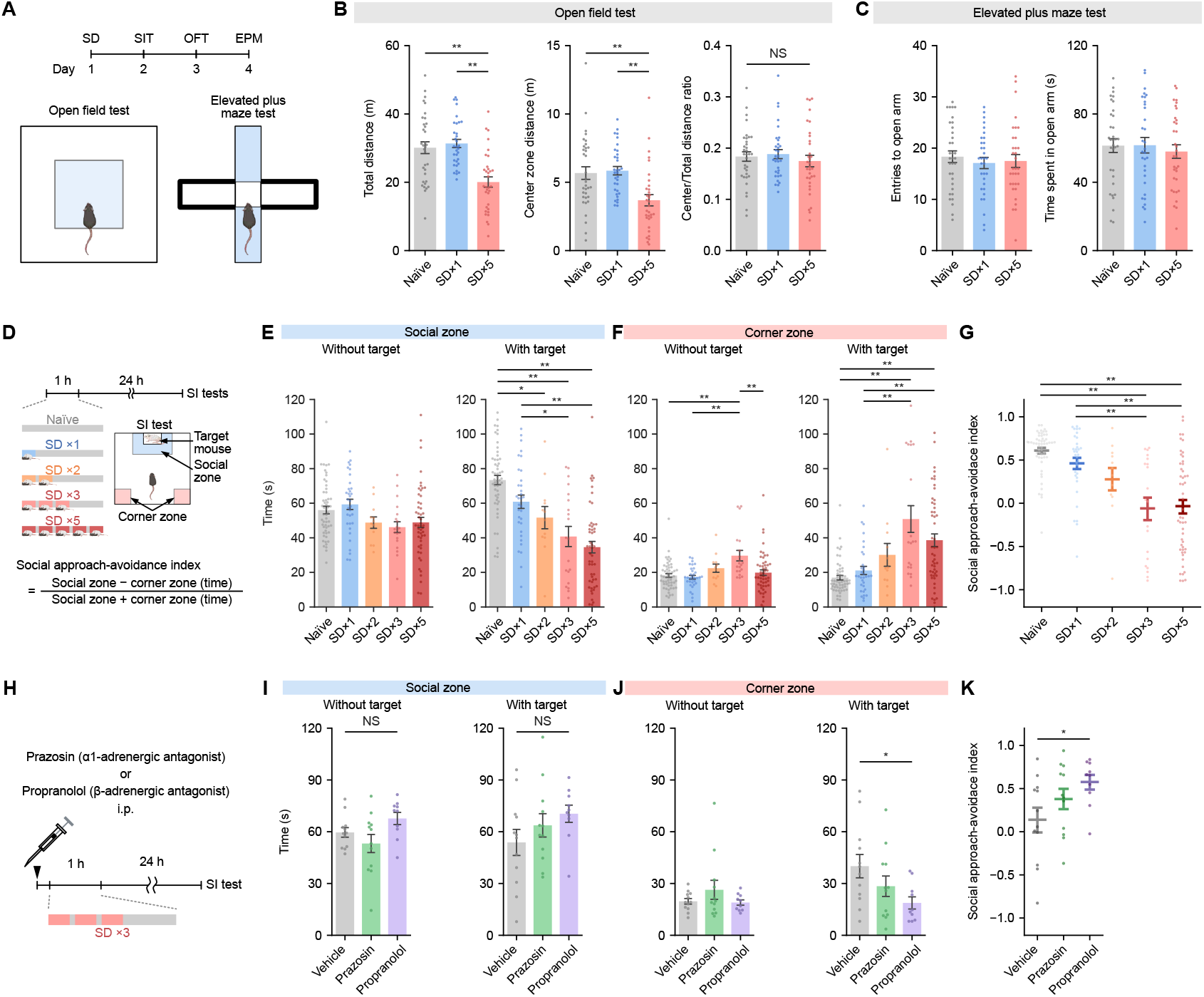
Behavioral alteration by acute social defeat stress and its regulation mediated by adrenergic receptors. (**A**) Schematic illustrating behavioral experiments. The mice used in Figure 2A–2D were subjected to the open field test and elevated plus maze test. (**B**) Quantification of the total ambulation distance (left), ambulation distance in the center zone (middle), and ratio of total and center zone ambulation distances (right) in the open field test. The ratio of ambulation distance in the center zone and total ambulation distance was used to normalize the difference in total ambulation distances. Naïve, n = 34; SD×1, n = 33; SD×5, n = 33. NS, not significant; **P < 0.01. (**C**) Quantification of the number of entries to open arm (let) and the time spent in the open arm (left) in the elevated plus maze test. Naïve, n = 34; SD×1, n = 32; SD×5, n = 33. One mouse in the SD×1 group fell from the maze and was excluded from analysis. (**D**) Schematic illustrating behavioral experiments for determining subthreshold levels of social defeat stress. For SD×2 and SD×3, different duration of social defeat and sensory contact with a clear divider were employed (see Materials and Methods for details). (**E, F**) Quantification of time spent in the social and corner zones with or without a target mouse. Data include the mice used in Fig. 2A–2D. Naïve, n = 54; SD×1, n = 33; SD×2, n = 12; SD×3, n = 20; SD×5, n = 51. *P < 0.05, **P < 0.01, one-way analysis of variance (ANOVA) followed by Tukeys test. (**G**) Quantification of the social approach avoidance index. **P < 0.01, one-way ANOVA followed by Holm-Sidak test. (**H**) Schematic illustrating behavioral experiments for pharmacological inhibition of adrenergic receptors. C57BL/6J mice were subjected to SD×3 at 30 min after intraperitoneal administration of saline, 1 mg/kg prazosin, or 10 mg/kg propranolol. (**I, J**) Quantification of time spent in the social and corner zones with or without a target mouse. Vehicle, n = 12; prazosin, n = 12; propranolol, n = 10. NS, not significant; *P < 0.05, one-way ANOVA followed by Dunnetts test. (**K**) Quantification of the social approach avoidance index. *P < 0.05, one-way ANOVA followed by Dunnetts test. Data are represented as mean *±* SEM. The illustrations of mice in (A, D) were created with BioRender.

**Fig. S3.**
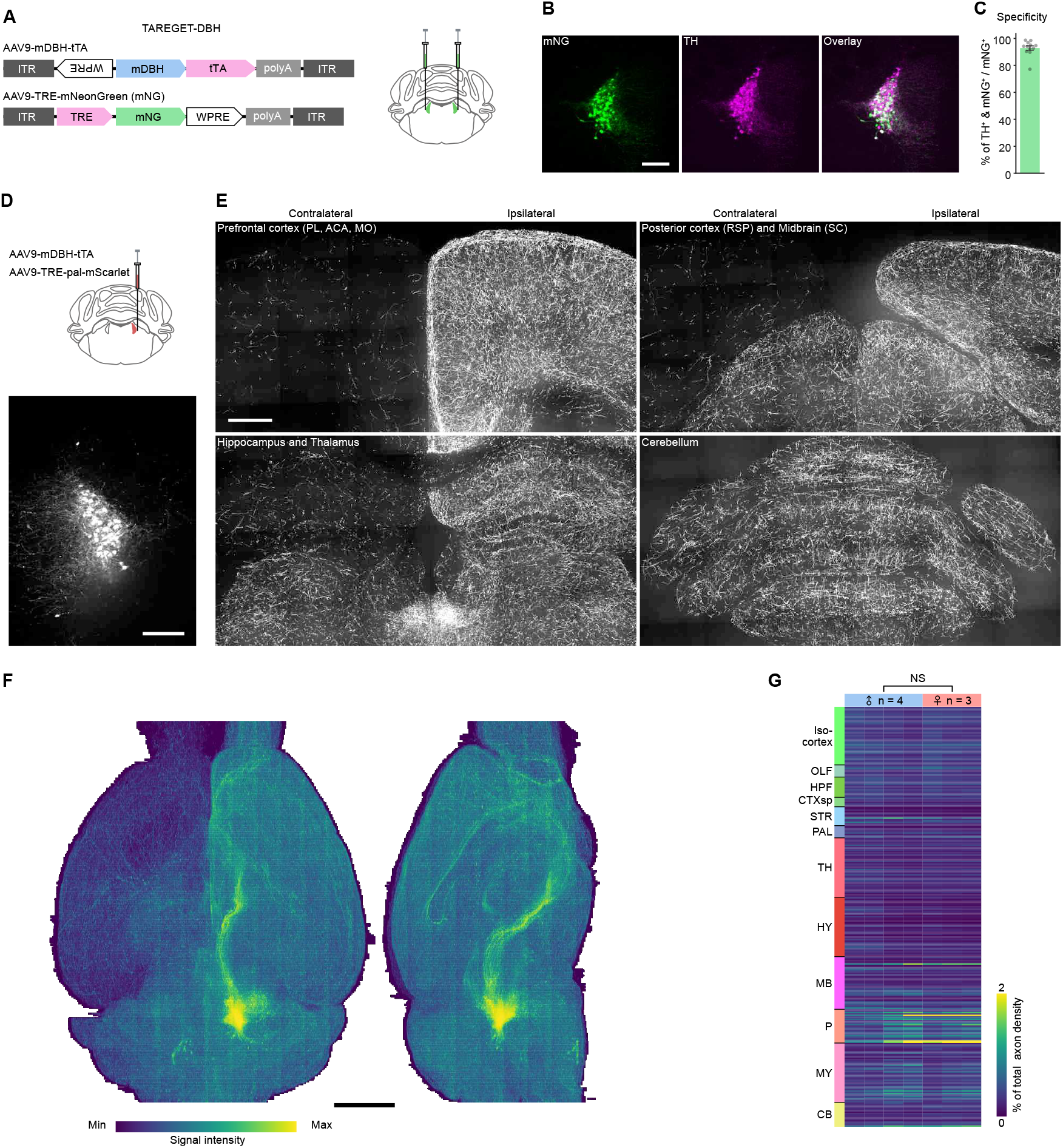
AAV-mediated LC^NA^ neuron-selective gene expression for brain-wide axonal mapping. **(A)** Schematic illustrating the AAV vectors for NA neuron-selective gene expression, named TAREGET-DBH. AAV9-mDBH-tTA includes the inverted WPRE at the 3 upstream of mouse DBH promoter. (**B**) Representative images of the mNG expression, TH immunofluorescence, and their overlay. Scale bar, 200 µm. (**C**) Specificity of mNG expression controlled by mDBH promoter-driven tTA. The percentage of mNG-positive cells that also expressed TH is indicated. n = 12 injection sites in six mice. (**D**) Schematic illustrating experiments for brain-wide axonal mapping and a representative image of LC^NA^ neurons expressing membrane-targeted fluorescent protein, pal-mScarlet. Scale bar, 200 µm. (**E**) Representative images of axons labeled with pal-mScarlet. PL, prelimbic area; ACA, anterior cingulate area; MO, somatomotor area; RSP, retrosplenial area; SC, superior colliculus. Scale bar, 500 µm. (**F**) Representative 3D reconstruction of the whole brain in a mouse injected with AAV9-mDBH-tTA and AAV9-TRE-pal-mScarlet. Scale bar, 2 mm. (**G**) Heatmap of relative axon density, adjusted based on the volume of each brain region and the total label densities across 311 regions in individual male and female brains. OLF, olfactory areas; HPF, hippocampal formation; CTXsp, cortical subplate; STR, striatum; PAL, pallidum; TH, thalamus; HY, hypothalamus; MB, midbrain; P, pons; MY, medulla; CB, cerebellum.

**Fig. S4.**
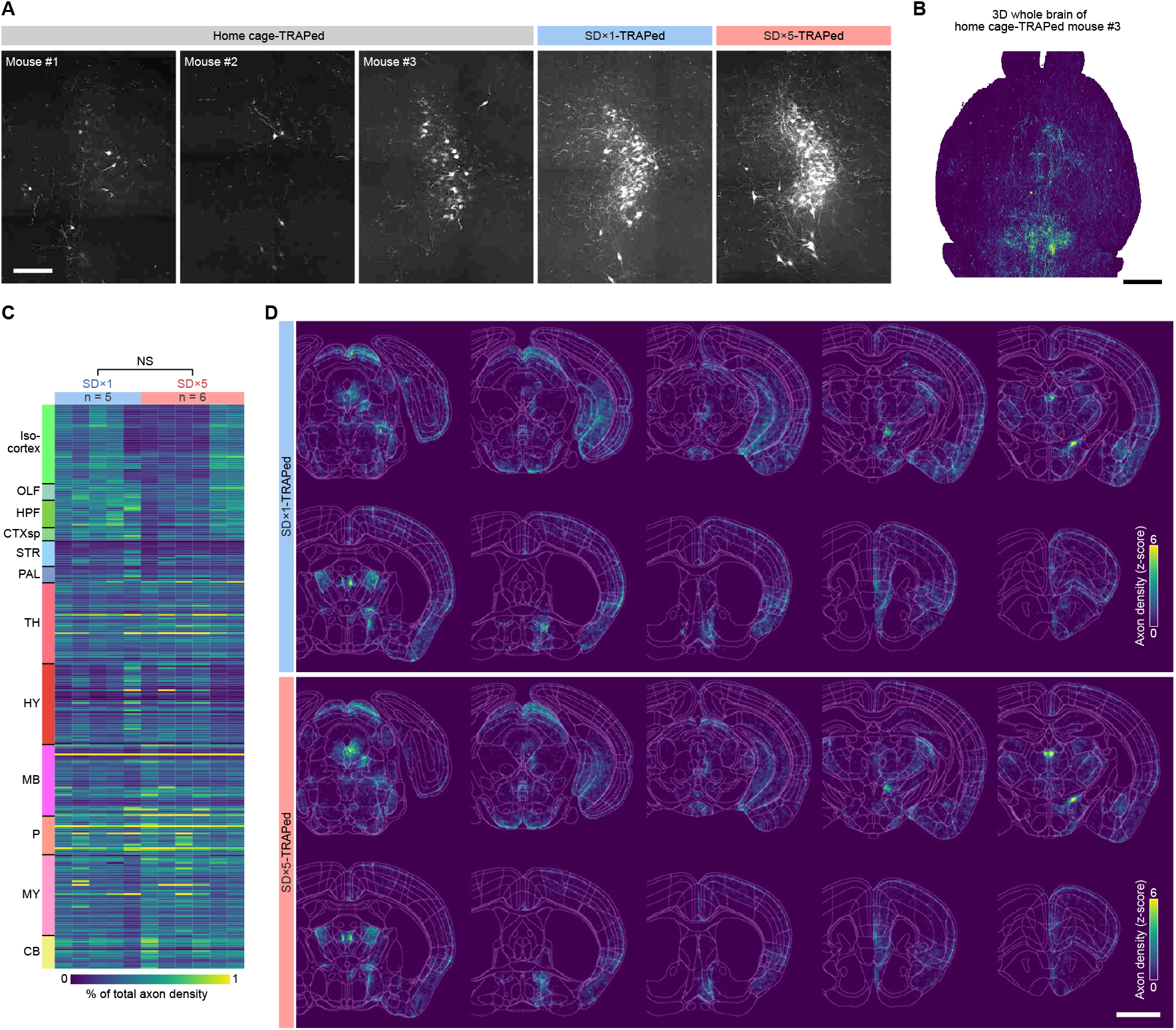
Mesoscale projectome of stress-responsive LC^NA^ neurons. (**A**) Representative images of the LC in TRAP2 mice subjected to home cage condition, SD×1, or SD×5. Scale bar, 200 µm. (**B**) Representative 3D reconstruction of the whole brain in a TRAP2 mouse subjected to home cage conditions, mouse 3 represented in (A). Scale bar, 2 mm. (**C**) Heatmap of relative axon density from individual mice, with adjustment based on the volume of each brain region and the total label densities across 311 regions in individual brains. OLF, olfactory areas; HPF, hippocampal formation; CTXsp, cortical subplate; STR, striatum; PAL, pallidum; TH, thalamus; HY, hypothalamus; MB, midbrain; P, pons; MY, medulla; CB, cerebellum. NS, not significant. (**D**) Coronal views of the voxel heatmap of mean z-score-normalized axonal densities for TRAP2 mice subjected to SD×1 and SD×5. Scale bar, 2 mm.

**Fig. S5.**
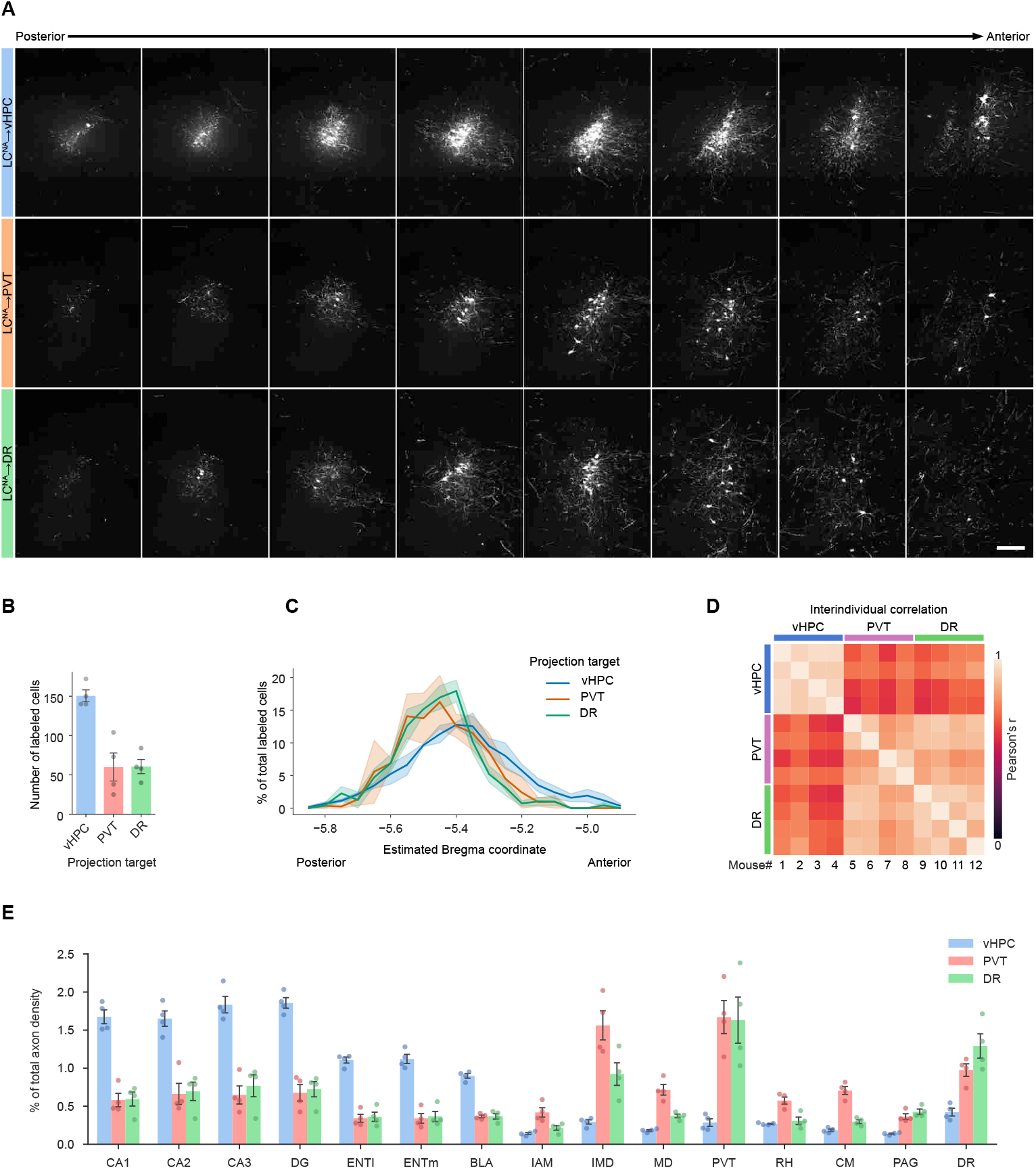
Anatomical characteristics of the LC^NA^→vHPC, LC^NA^→PVT, and LC^NA^→DR neurons. **(A)** Representative serial section images of the LC^NA^ neurons retrogradely targeted from the vHPC, PVT, or DR. Maximum intensity projection images were represented at 50-µm intervals, each representing a 50-µm-thick section along the anteroposterior axis. Scale bar, 200 µm. The number of pal-mScarlet-labeled cells in the LC retrogradely targeted from the vHPC, PVT, or DR. (**C**) Anteroposterior distribution of pal-mScarlet-labeled cells in the LC retrogradely targeted from the vHPC, PVT, or DR. Percentages of labeled cells at each bregma coordinate relative to the total number of labeled cells were plotted. (**D**) Pearson correlation matrix of interindividual axonal projection patterns in 12 mice (four mice per projection-defined group). Pairwise correlation coefficients (Pearsons r) were computed for relative axon densities, adjusted based on the volume of each brain region and the total label densities across 311 regions. (**E**) Quantification of the relative axon densities in 15 representative brain regions. CA1, hippocampal field CA1; CA2, hippocampal field CA2; CA3, hippocampal field CA3; DG, dentate gyrus; ENTl, entorhinal area, lateral part; ENTm, entorhinal area, medial part; BLA, basolateral amygdalar nucleus; IAM, interanteromedial nucleus of the thalamus; IMD, intermediodorsal nucleus of the thalamus; MD, mediodorsal nucleus of the thalamus; RH, rhomboid nucleus; CM, central medial nucleus of the thalamus. Data are represented as mean *±* SEM.

**Fig. S6.**
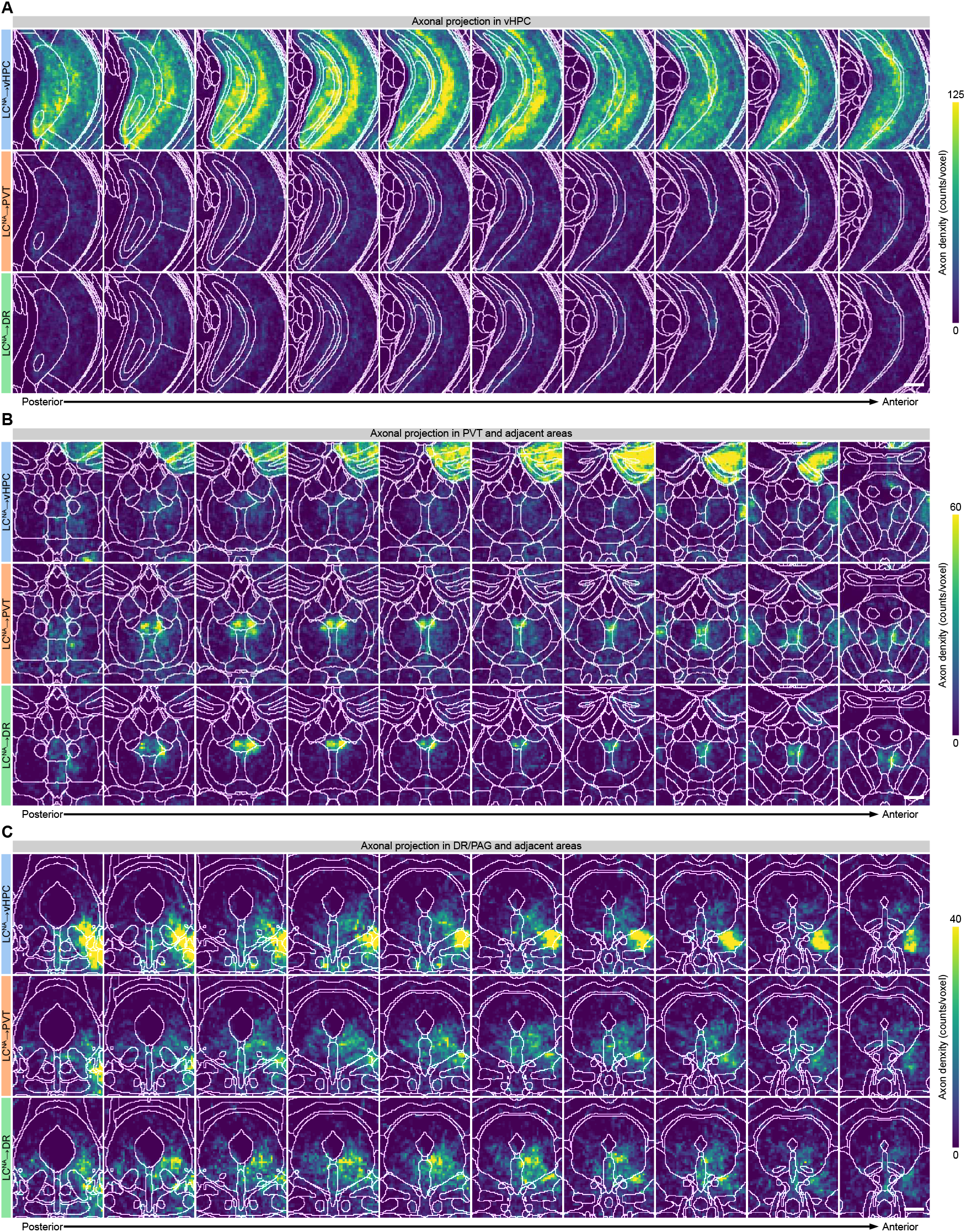
Distinct axonal distribution patterns of anatomically defined LC^NA^ subpopulations in the main target brain regions, vHPC, PVT, and DR. (**A–C**) Magnified coronal views of the voxel heatmap of mean axonal densities in brain regions including the vHPC (A), PVT (B), and DR (C) for retrogradely targeted LC^NA^→vHPC (top), LC^NA^→PVT (middle), and LC^NA^→DR (bottom) neurons. Serial section images were presented at 50-µm intervals. Scale bars, 500 µm

**Fig. S7.**
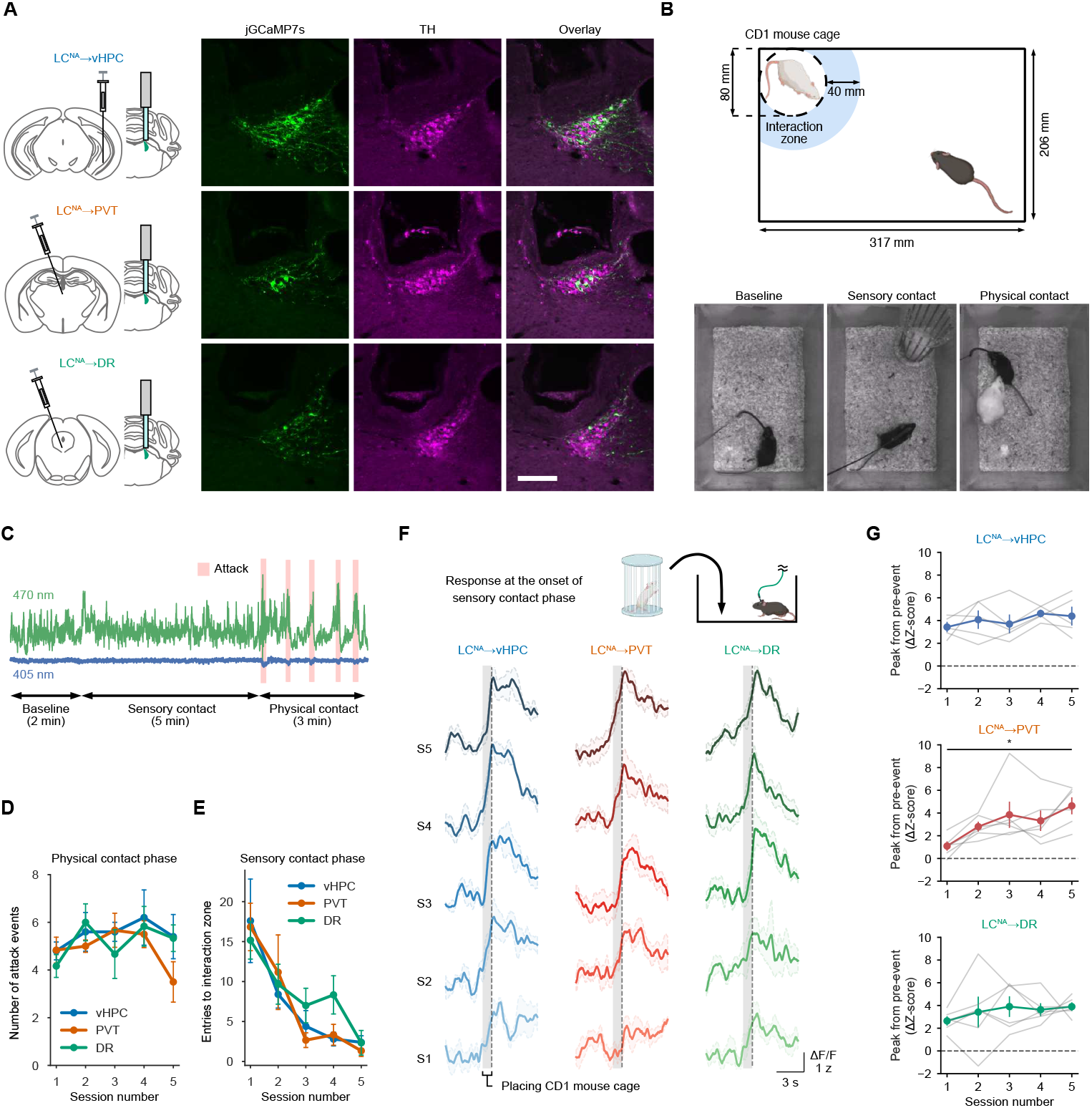
Distinct neural response dynamics in fiber photometry recordings of projection target-defined LC^NA^ neurons. (**A**) Representative images of the jGCaMP7s-expressing cells and TH-immunostained cells in the LC retrogradely targeted from the vHPC, PVT, or DR using AAVrg-hSyn-iCre. Scale bar, 200 µm. (**B**) Schematic illustrating the sensory contact phase (top) and representative video frames for baseline, sensory contact, and physical contact phases during a social defeat stress session for fiber photometry recording (bottom). The onset of social approach was defined as the entry of the body center into the interaction zone, except for repeated entries within 1 s. (**C**) Representative trace of recording during the first stress session from jGCaMP7s-expressing LC^NA^ neurons retrogradely targeted from the vHPC. Calcium-dependent signals (green) excited by 470 nm light and isosbestic signals (blue) excited by 405 nm light are indicated. The timeframes during attacks are highlighted in red. (**D**) Quantification of the number of attack events by aggressor mice in the physical contact phase across sessions. (**E**) Quantification of the number of entries to the interaction zone in the sensory contact phase across sessions. (**F**) Neuronal activities of the LC^NA^→vHPC, LC^NA^→PVT, and LC^NA^→DR neurons at the onset of sensory contact phase (insertion of the CD1 mouse cage) across sessions (S1S5). Peri-event plot showing mean neuronal activity 5 s before and after insertion of the social grid cage. The grid cage was manually inserted into the animal housing cage within approximately 1 s, and the 1-s insertion period is highlighted in gray. (**G**) Quantification of peak response amplitudes after the event onset (0 to 2 s) from pre-event baseline. *P < 0.05, one-way repeated measures ANOVA followed by Holm-Sidak test. Data are represented as mean *±* SEM. The illustrations of mice in (B, F) were created with BioRender.

**Fig. S8.**
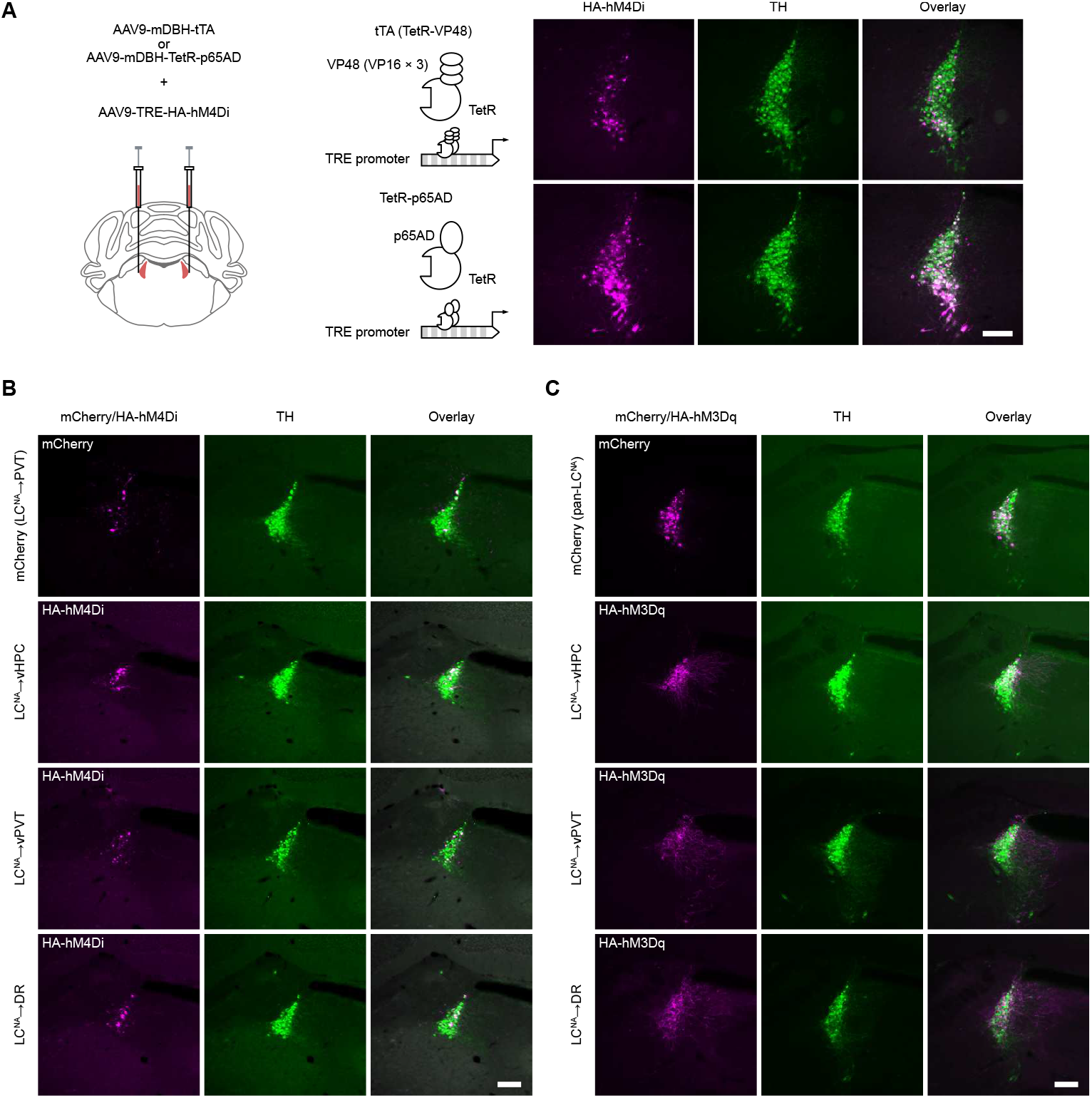
DREADD receptor expression in anatomically defined LC^NA^ neurons. (**A**) Schematic illustrating the strategy for inducing HA-hM4Di expression in LC^NA^ neurons (left). Representative images of HA-hM4Di-expressing cells and TH-immunostained cells in the LC are shown (right). HA-hM4Di was immunostained for HA tag. (**B**) Representative images of the LC in the projection target-selective chemogenetic inhibition experiment. HA-hM4Di and mCherry expressions were driven by Tet-p65AD. (**C**) Representative images of the LC in the projection target-selective chemogenetic activation experiment. HA-hM3Dq and mCherry expressions were driven by conventional tTA. Scale bars, 200 µm.

**Fig. S9.**
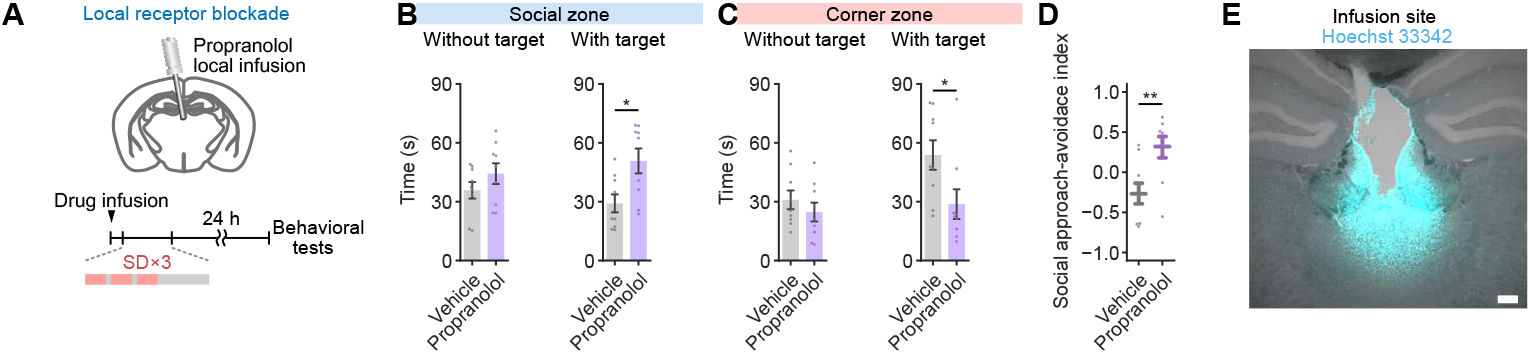
Involvement of the PVT beta-adrenergic receptor in social avoidance learning. (**A**) Schematic illustrating behavioral experiments for local pharmacological blockade of beta-adrenergic receptors in the PVT. Mice were subjected to SD×3 episodes 15 min after administration of 1 µg propranolol into the PVT via an implanted cannula. (**B, C**) Quantification of time spent in the social (B) and corner (C) zones with or without a target mouse. Vehicle, n = 9; propranolol, n = 9. *P < 0.05, Students t-test. (**D**) Quantification of the social approach avoidance index. (**E**) Representative image of the PVT in mice that received local infusions of propranolol. After behavioral experiments, mice received local infusion of a nuclear dye, Hoechst 33342 to visualize the infusion site. Scale bars, 200 µm. Data are represented as mean *±* SEM.

